# Integrated GWAS and transcriptome analyses reveal candidate genes associated with betalain-related pigmentation under heat stress in *Amaranthus tricolor*

**DOI:** 10.64898/2026.10.05.756862

**Authors:** Ya-Ping Lin, Hina Okubo, Akito Nashiki, Yosuke Yoshioka, Yuan-Yun Zhang, Hieng-Ming Ting, Sachiko Isobe, Kenta Shirasawa, Ken Hoshikawa

**Affiliations:** World Vegetable Center, P.O. Box 42, Shanhua, Tainan 74199, Taiwan; Graduate School of Science and Technology, University of Tsukuba, Tennodai 1-1-1, Tsukuba 305-8572, Ibaraki, Japan; Institute of Life and Environmental Sciences, University of Tsukuba, Tennodai 1-1-1, Tsukuba 305-8572, Ibaraki, Japan; Institute of Plant Biology, National Taiwan University, Taipei 10617, Taiwan; Kazusa DNA Research Institute, Kazusa-kamatari, Kisarazu, Chiba 292-0818, Japan; Biological Resources and Post-harvest Division, Japan International Research Center for Agricultural Sciences, Ohwashi 1-1, Tsukuba, Ibaraki, 305-8686, Japan

**Keywords:** *Amaranthus tricolor*, orphan crop, heat stress tolerance, betalain biosynthesis, GWAS

## Abstract

*Amaranthus tricolor* is a climate-resilient orphan crop whose betalain pigments contribute to nutritional quality and stress tolerance. However, breeding efforts have traditionally relied on field yield trials due to the limited availability of genomic information and genetic resources. To address this gap, we first used the Illumina and PacBio sequencing platforms to assemble a novel reference genome of *A. tricolor*, using the Japanese cultivar Bayam. By combining this reference genome with a core collection developed in our previous study, we aimed to identify genomic regions associated with pigmentation-related traits across winter and summer seasonal environments. Betalains are antioxidant phytonutrients that contribute to abiotic stress tolerance, making the regulation of their biosynthesis under heat stress particularly relevant for developing climate-resilient crops such as *A. tricolor.* As a result, ATR1.0CH03_5131103, which is associated with *ATR1.0ch03g000565,* was identified under the winter seasonal environment. Meanwhile, transcriptomic profiles in this study revealed that *ATR1.0ch03g000565* was differentially expressed under both control and heat stress conditions, suggesting that it may be associated with the regulation of *CYP76AD1* expression under optimal conditions. Under the elevated-temperature treatment, transcript levels of *CYP76AD1* and *5GT* decreased in Am087, coinciding with lower betacyanin_dominated absorbance indices. *cDOPA*-like homologs were upregulated; however, pigmentation-related absorbance remained reduced. Further molecular experiments, such as CRISPR-Cas9 knockouts or overexpression studies, are necessary to validate the regulatory mechanisms of *CYP76AD1* and *5GT* under heat stress.

## Introduction

Malnutrition has long been a critical threat to human health, with an estimated 821 million people in South Asia and Africa suffering from inadequate food consumption and nutritional deficiencies (FAO, 2018). Orphan crops, which include various indigenous or local vegetables in developing countries, are increasingly recognized as valuable resources for establishing sustainable food systems that can provide nutritious foods in an environmentally resilient manner (Sarker et al., 2022). One of the indigenous vegetables, *Amaranthus*, is well-adapted to a range of abiotic stresses, including drought, salinity, heat, and heavy metal exposure (Sarker et al., 2022; Reyes-Rosales et al., 2023). The genus *Amaranthus* comprises approximately 50□70 species globally, some of which are consumed as pseudo-grains, others as leafy vegetables. *Amaranthus tricolor* has traditionally been cultivated in Africa and Asia, where they are valued as low-input, nutrient-rich, and medicinal vegetables (Singh et al., 2024). Their high nutrient content and tolerance to abiotic stresses make leafy amaranths promising candidates for climate-resilient agriculture.

Most breeding efforts in *A. tricolor* have focused on improving yield, with breeders selecting relatively stable and high-yield lines based on field trials. Recently, advances in high-throughput sequencing technologies have accelerated the study of functional genes in *A. tricolor*. For instance, transcriptomic analyses have identified *AmCYP76AD1* as a crucial gene in betalain biosynthesis, providing a pathway for improving betalain content through marker-assisted selection or further functional studies (Chang et al., 2021). A current study used a selected-genome sequencing approach to develop single-nucleotide polymorphisms (SNPs) for a worldwide *A. tricolor* collection and to develop a core collection (Hoshikawa et al., 2023). Given that *A. tricolor* is genetically distant from *A. hypochondriacus*, the first *Amaranthus* species with a reference genome, and possesses a different chromosome number, mapping rates to the *A. hypochondriacus* reference genome are relatively low, consequently reducing the SNP number (Nguyen et al., 2019; Lin et al., 2021). A newly assembled *A. tricolor* genome, particularly from a market-dominated variety, would provide a species-specific reference that significantly improves SNP marker development and functional gene studies.

Betalains are unique pigments found in most species within the order *Caryophyllales*, including the family *Amaranthaceae* (Sarker et al., 2022). As phytonutrients, betalains have pharmaceutical applications, with intermediate products in the betalain pathway used to treat Parkinson’s disease (Galanie et al., 2015). Additionally, betalains are natural colorants widely used in the food industry (Galanie et al., 2015). In plants, betalains have diverse functions, including attracting pollinators, repelling herbivores, protecting tissues from ultraviolet radiation, and removing reactive oxygen species (ROS), all of which contribute to abiotic stress tolerance (Albano et al., 2015). Betalains are classified as betacyanins and betaxanthins, which are red and yellow, respectively. Betacyanins include four types of pigments, namely, amaranthin, betanin, gomphrenin, and decarboxybetanin (Sarker et al., 2022). The shikimate pathway supplies l-tylosin, a precursor of betalain biosynthesis (Liu et al., 2019). The betalain biosynthesis pathway involves five key enzymes, including cytochrome P450 76AD1 (*CYP76AD1*: *LOC130803140*), L-dihydroxyphenylalanine 4,5-dioxygenase (*DODA*: *LOC130803136*), cyclo-DOPA 5-O-glucosyltransferase (*cDOPA*: *LOC130809389*), betanidin-5-O-glucosyltransferase (*5GT*: *LOC130825231*), and tyrosine decarboxylase (*TyDC*: *LOC130817294*) (Chang et al., 2021; Figure 1). *CYP76AD1* serves as the first enzyme to initiate the betalain biosynthesis, which is followed by either through *DODA* and *5GT* or *CYP76AD1* and *cDOPA* to produce betanin, and via *DODA* to synthesize betaxanthin.

**Figure 1.**
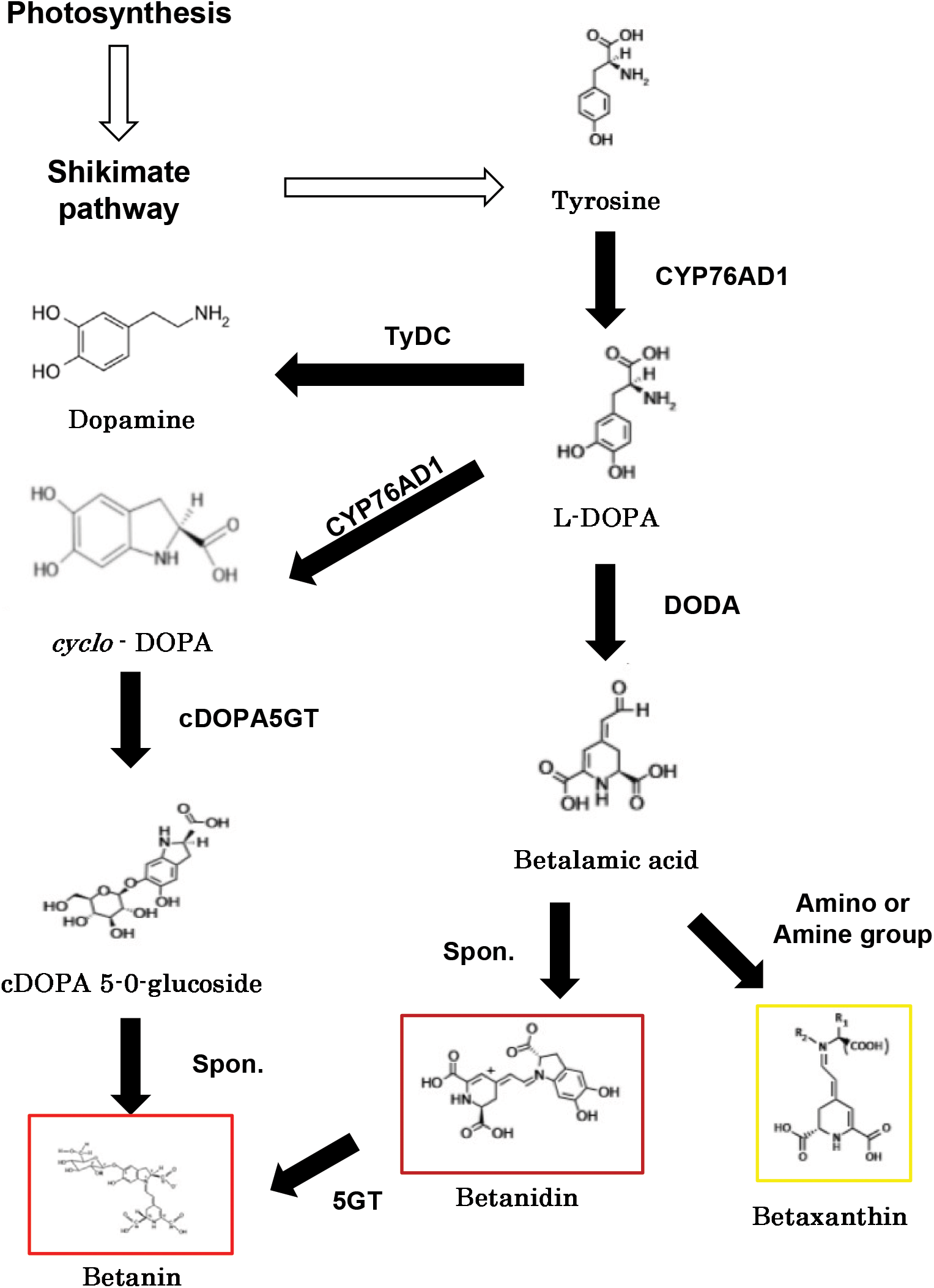
Pathway of betalain biosynthesis. This figure illustrates the betalain biosynthesis pathway, highlighting key enzymes involved in betacyanin and betaxanthin synthesis. The pathway includes *CYP76AD1* (*cytochrome P450 76AD1*), L-DOPA (L-3,4-dihydroxyphenylalanine), *DODA* (*L-dihydroxyphenylalanine 4,5-dioxygenase*), *cDOPA5GT* (*cyclo-DOPA 5-O-glucosyltransferase*), *B5GT*, (*betanidin-5-O-glucosyltransferase*), and *TyDC* (*tyrosine decarboxylase*) with arrows indicating metabolic conversions. The squares in the figure indicate the colors of each pigment. The figure was adapted based on experimental findings from this study and relevant literature sources (Chang et al., 2021, Figure 1e).

**Figure 2.**
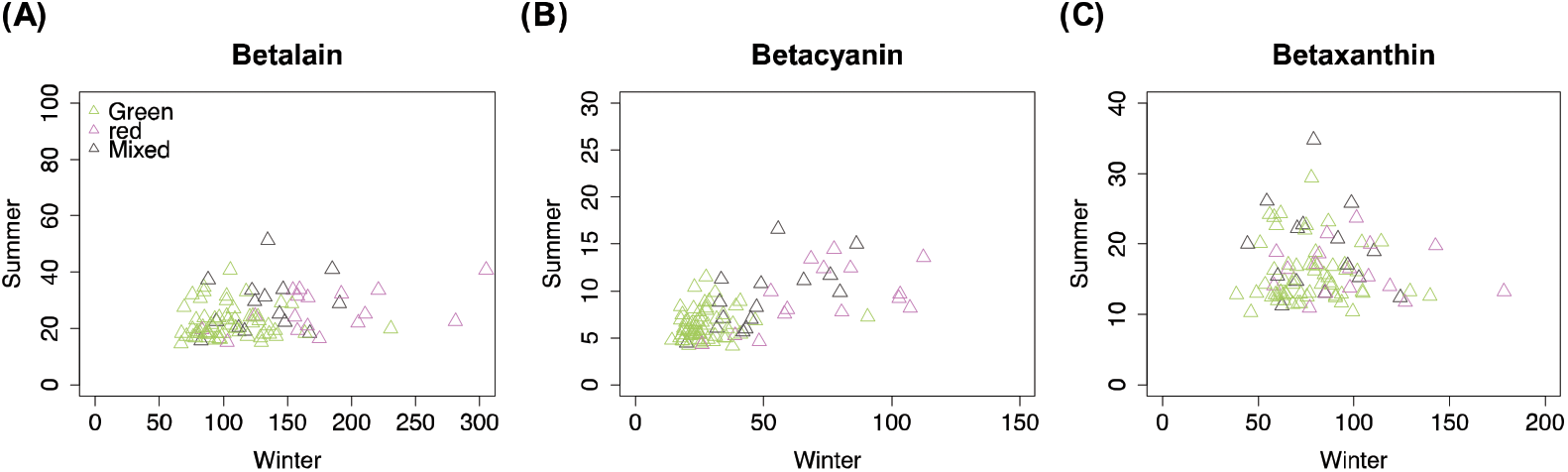
Seasonal comparison of spectrophotometric betalain absorbance indices. (A) Betalain absorbance index (BAI = A538 + A480); (B) index-A538 (betacyanin□dominated absorbance); (C) index-A480 (betaxanthin□dominated absorbance). The x- and y-axes represent absorbance values measured in winter and summer, respectively. Each triangle represents one accession (n = 3 per accession per season). Green, violet, and grey indicate green, red, and mixed color amaranths, respectively.

Additional transcription factors are involved in the regulation of betalain biosynthesis. Previous studies have highlighted the WRKY family members, which were not only abundant in the amaranth transcriptomic profile but also demonstrated the ability to bind to the *CYP76AD1* promoter, thereby promoting betacyanin synthesis (Yang et al., 2024). Overexpression of *AtrWRKY42-2* has been shown to upregulate *CYP76AD1*, *B5-GT*, *B6-GT*, and *TyDC* (Yang et al., 2024). Similar findings were reported in pitaya (*Hylocereus monacanthus*), underscoring the role of WRKY transcription factors in betacyanin biosynthesis (Zhang et al., 2020). Moreover, the MYB family has also been implicated in the regulation of betacyanin biosynthesis. In pitaya, *HuMYB132* enhances the transcriptional activity of *HuCYP76AD1–1* and *HuDODA1* by binding to their promoters (Xie et al., 2023). Silencing *HuMYB132* leads to reduced betalain accumulation and downregulation of betalain biosynthetic genes. In pear, *PyWRKY26* has been found to target *PyMYB114*, thereby regulating anthocyanin accumulation (Li et al., 2020). These findings suggest a complex, yet relatively unexplored, regulatory network governing betalain biosynthesis.

Although *A. tricolor* is gaining attention for its resilience to abiotic stress and high nutritional value, breeding efforts have been constrained by limited genomic resources and insufficient integration of functional genetic variation into improvement programs. Recent advances in sequencing technologies have facilitated the identification of genes involved in betalain biosynthesis. However, effective strategies for enriching, characterizing, and deploying betalain-associated genetic resources in breeding programs remain largely unexplored. The lack of a high-quality, species-specific reference genome has further limited the translation of molecular insights into practical breeding applications. To address these challenges, we assembled a novel reference genome from an elite Japanese cultivar, ‘Bayam’, to enable high-resolution genomic analyses. Using this reference, we generated an improved SNP dataset for a core germplasm collection and conducted a genome-wide association study (GWAS) to identify tentative marker–trait associations for pigmentation-related traits across winter and summer growing seasons. In parallel, transcriptomic analyses were employed to 1) refine GWAS candidate regions, 2) to evaluate the expression of key betalain biosynthesis genes (*5GT*, *cDOPA*, *CYP76AD1*, *DODA*, and *TyDC*) under control and heat stress conditions, and 3) to identify additional candidate genes with potential utility for marker-assisted selection and the development of climate-resilient *A. tricolor* cultivars.

## Materials and Methods

### Plant Material

This study used the core collection of *Amaranthus tricolor* established in a previous study (Hoshikawa et al., 2023; Supplementary Table 1) to examine betalain-related absorbance across winter and summer seasons in Taiwan. The leaf color was defined by color rate: Green (G) indicates all green without red color; Mix (M) stands for 10-50% red mixed with green color; Mix Red (MR) refers to >50 % red color; Red (R) indicates almost red color on leaves. A total of 95 lines, categorized into 60 green, 19 red, and 15 mixed-color variants, were cultivated in a greenhouse for phenotypic assessment. *A. tricolor* cv. Bayam was selected as a reference cultivar because of its genetic uniformity, derived from seven generations of selfing, and its relatively stable betalain-related absorbance across seasonal environments. The winter cultivation (October 2021) recorded daily max/min/mean temperatures of 31.2/23.1/26.7□°C, and the summer cultivation (August 2022) recorded 33.8/25.1/28.5□°C. We contrast winter vs summer environmental conditions (greenhouse means 26.7□°C vs 28.5□°C); given concomitant changes in irradiance, vapor pressure deficit (VPD), humidity, and photoperiod, we avoid interpreting this contrast as a controlled heat□stress treatment. Plants were grown for one month, after which seedlings were harvested for betalain-related absorbance analysis. For each experimental condition, three biological replicates were used, with each replicate consisting of pooled leaf samples from three individual plants.

### Measurement of betalain-related absorbance

Betalain-related absorbance was quantified following the protocol outlined by Ravichandran et al. (2013). Samples were flash□frozen in liquid nitrogen, freeze□ried, and finely pulverized with a sterile pestle and mortar. For extraction, 0.10 g of powdered material was extracted in 10 mL 50% (v/v) aqueous ethanol by vortexing followed by brief sonication, and then centrifuged at 6000 rpm for 10 min. The supernatant was collected and recentrifuged to remove residual particulates. UV-Vis spectra were recorded using 1□cm path□length cuvettes. The extraction volume (10□mL) and spectrophotometric path length (1□cm) were kept consistent across all experiments. We quantified betacyanin□dominated absorbance (indices-A538) and betaxanthin□dominated absorbance (indices-A480) as relative indicators of pigment abundance rather than as estimates of absolute concentrations. A combined betalain absorbance index was calculated as the sum of absorbance values at 538 nm and 480 nm. Because ethanolic extracts may contain co-extracted chlorophylls and carotenoids, which can contribute to absorbance, particularly near 480 nm and to a lesser extent near 538 nm, these measurements are interpreted as relative indices. Each line was evaluated in triplicate (three biological replicates). To assess the seasonal variation in betalain-related absorbance, an ANOVA test was conducted, treating season and plant color as fixed effects.

### Genome assembly of *Amaranthus tricolor* cv. Bayam

The selfing F_7_ generation of *A. tricolor* cv. Bayam was used as the model cultivar for *A. tricolor* to construct a novel reference genome. Illumina and PacBio sequencing systems were used to assemble a new chromosome-level reference genome. For Illumina sequencing, genomic DNA was extracted from young *A. tricolor* cv. Bayam leaves provided by NOGUCHI Seed Co. LTD, with a Genomic-Tip (Qiagen, Hilden, Germany). A short-read sequencing library was constructed using the TruSeq DNA polymerase chain reaction-Free LT Sample Prep Kit (Illumina, San Diego, CA, USA) and sequenced using the Illumina HiSeq X platform. K-mer frequency was used to estimate the genome size using jellyfish (Marçais and Kingsford, 2011). For PacBio sequencing, DNA libraries were constructed using the SMRTbell Express Template Prep Kit 2.0 (PacBio, Menlo Park, CA, USA) and sequenced for six Single-molecule real-time (SMRT) cells (1M v3) on the PacBio platform. The reads were assembled with Falcon Unzip (https://github.com/PacificBiosciences/FALCON/) to obtain primary contigs and haplotigs of the genome. Potential sequence errors in the assembled sequences were corrected twice using long reads with Arrow implemented in the SMRT Link (PacBio). The pseudomolecular sequences were constructed based on a genetic map developed from the F_2_ population (n = 380). The F_2_ population was derived from a cross between Bayam and Asian Red, and SNP calling followed the procedures described by Shirasawa et al. (2016). Briefly, genomic DNA was extracted from the leaves and digested with two restriction enzymes, *Pst*I and *Msp*I. Libraries were sequenced on a DNBSEQ-G400 platform (MGI). The sequencing reads were mapped onto primary contigs using Bowtie2 (Langmead and Salzberg, 2012). High-confidence biallelic SNPs were identified using the mpileup option of SAMtools (Li et al., 2009) and filtered using VCFtools (Danecek et al., 2011) under the following conditions: read depth ≥ 5, SNP quality = 10, minor allele frequency ≥ 0.2, and proportion of missing data < 50%. A genetic map was constructed using Lep-Map3 (Rastas, 2017). The contig sequences were anchored to a genetic map to build the pseudomolecular sequence ALLMAPS (Tang et al., 2015). Repetitive sequences were identified using the RepeatMasker (Smit et al.). In accordance with RepeatMasker, repeat elements were classified into nine types: short interspersed nuclear elements, long interspersed nuclear elements, long terminal repeat elements, DNA elements, small RNA, satellites, simple repeats, low-complexity repeats, and unclassified elements.

### Genome Annotation

To comprehensively investigate gene distribution across the entire genome, we performed a thorough annotation, encompassing both coding and repetitive regions. This annotation was based on a combination of ab initio predictions, RNA, and protein evidence. For ab initio predictions, gene models were generated using AUGUSTUS v3.4.0 with the “Arabidopsis” parameter (Stanke et al., 2008). RNA evidence was derived from transcriptomic data obtained from *Amaranthus* species grown under controlled conditions. We weighted RNA/protein evidence heavily to reduce model bias; we now highlight this caveat and its potential impact on gene model fidelity. These RNA-Seq reads were aligned to the newly assembled reference genome using HISAT2 v2.2.1 (Kim et al., 2019), and transcripts were subsequently reconstructed with StringTie v2.2.2 (Pertea et al., 2015). Open Reading Frames (ORFs) longer than 100 amino acids were predicted using TransDecoder v5.7.0, referencing UniProt and Pfam domains (Haas et al., 2017). Protein evidence involved aligning sequences from *Amaranthus hypochondriacus*, *Amaranthus cruentus*, and *Spinacia oleracea* to the new reference genome using GenomeThreader v1.7.1 (Gremme et al., 2005). The ab initio predictions, RNA evidence, and protein evidence were integrated with respective weights of 1, 5, and 5 using EVidenceModeler v1.1.1 (Haas et al., 2008) to construct consensus gene models. Functional annotation of the predicted genes was performed using eggNOG-mapper v2.1.9 (Cantalapiedra et al., 2021). Additionally, homologous genes related to betalain biosynthesis, including *CYP76AD1* (*QOP57914.1*), *DODA* (*QOP57916.1*), *cDOPA* (*QOP57917.1*), *5GT* (*AJY59053.1*), and *TyDC* (*AJW81117.1*), were identified through BLAST with an *E-value* threshold of < 10^-50.

### Genome-Wide Association Study (GWAS) of betalain-related absorbance

We conducted a genome-wide association study (GWAS) to identify genetic variants linked to betalain-related absorbance. Raw reads generating using Illumina HiSeq X platform from double digest restriction-site associated DNA sequencing (ddRAD-seq) were retrieved from the DDBJ Sequence Read Archive database (accession numbers DRA012903 - DRA012912; Supplementary Table 2). The reads were quality-filtered using SolexaQA v3.1.7.1 (Cox et al., 2010) and aligned to the newly constructed reference genome using BWA v0.7.17 (Li and Durbin, 2009). Single nucleotide polymorphisms (SNPs) were called using GATK v4.2.3.0 (McKenna et al., 2010) and filtered based on a minor allele frequency threshold of < 0.05. Prior to GWAS analysis, leaf color phenotypes were encoded as dummy variables. Two models, Blink and FarmCPU, were employed with the correction of the first three principal components via the Genome Association and Prediction Integrated Tool (GAPIT) (Wang and Zhang, 2021) to identify quantitative trait loci (QTL). Both raw *p-values* and *Benjamini–Hochberg False Discovery Rate* (*adjusted p-values*) were reported. The significance was defined as *adjusted p-values* < 0.05. For each significant SNP, putative positional candidate genes were defined as the annotated gene containing the SNP, when applicable, together with the immediately adjacent annotated genes upstream and downstream. For intergenic SNPs, the nearest annotated genes on either side of the SNP were retained. Candidate selection was therefore based on genomic adjacency rather than a predefined physical-distance window. SNP density was visualized with the CMplot package in R (Yin et al., 2024). Linkage disequilibrium (LD) analyses were conducted using TASSEL version 5.2.89 (Bradbury et al., 2007). A complete pairwise LD matrix was generated among all SNP markers within each chromosome. To assess the extent of LD decay, two complementary approaches were employed. First, the relationship between physical distance and LD (measured as *R^2^*) was estimated using locally weighted scatterplot smoothing (LOWESS) to characterize the decay pattern across each chromosome, implemented with the lowess() function in R. Second, mean *R^2^* values were computed within non-overlapping 100-kb sliding windows to estimate local LD decay across the genome. The results from both approaches were visualized using plot() function in R. In the overall LD decay, the purple curve is the LOWESS curve; the blue curve indicates the mean of sliding windows LD.

### RNA sequencing under heat stress and control conditions

Seeds of *Amaranthus* accessions Bayam, Am087, and Am127 were sown in plastic trays filled with a Nippi Gardening Soil No. 1 (Nihon Hiryo Co., Ltd.) and grown in a controlled growth chamber at 25°C with a photoperiod of 16 hours light and 8 hours dark. Plants were top□watered to container capacity daily, and this was applied at sowing. After one week, seedlings were either maintained at 25/25□C or subjected to a 35/25□°C regime (16□h/8□h light/dark); RH was not controlled, so temperature effects may have co□varied with VPD. Genotypes were randomized each week across benches to minimize positional effects. Leaves from one-month-old seedlings were harvested for RNA sequencing, with three biological replicates per accession under each treatment. Each biological replicate comprised leaves from three individual plants. RNA extraction and sequencing were performed by Rhelixa (Tokyo, Japan). Sequencing was performed using the Illumina HiSeq X platform. Details of the sequencing samples, the treatments, and sequencing coverage are provided in Supplementary Table 3.

### Identification of Differentially Expressed Genes (DEGs)

RNA sequencing reads were processed to remove low-quality sequences using Trimmomatic v0.39 (Bolger et al., 2014) and then aligned to the new reference genome using HISAT2 v2.2.1 (Kim et al., 2019). Alignment quality was assessed using samtools flagstat (Li et al., 2009). Estimated sequencing depth was calculated as (number of read pairs × 2 × 100 bp)/reference genome size. Gene-level read-assignment quality was assessed using the featureCounts summary statistics in Rsubreads for the 18 libraries included in the final analysis (Liao et al., 2019). Gene-wise dispersion estimates and the fitted mean–dispersion relationship were examined as model diagnostics. No additional low-count prefiltering was applied for this original within-accession DEG analysis.

For sample-level quality assessment, count data were variance-stabilized using the DESeq2 variance-stabilizing transformation (VST). Principal component analysis (PCA) was performed using the 500 most variable genes and visualized with the ggplot2 package in R (Wickham 2016). Sample-to-sample Pearson correlation coefficients were calculated from the VST-transformed expression matrix and visualized as a correlation heatmap. DEGs between elevated-temperature and control conditions within each accession were identified using DESeq2 (Love et al., 2014), with the design formula ∼ treatment. Statistical significance was defined as a *Benjamini–Hochberg adjusted P value* < 0.05.

To formally evaluate whether transcriptional responses to the temperature treatment differed among accessions, an additional genotype × treatment interaction analysis was conducted. For this analysis, genes with at least 10 counts in at least three libraries were retained before model fitting. DESeq2 was fitted using the factorial design ∼ genotype + treatment + genotype:treatment, with Bayam and the control treatment as reference levels. The genotype × treatment interaction test was performed using a likelihood-ratio test comparing the full model with a reduced model containing genotype and treatment main effects only. Pairwise interaction contrasts were additionally used to compare the treatment response of Am087 and Am127 with that of Bayam. Adjusted *P* values < 0.05 were considered significant. Because each genotype × treatment combination included three biological replicates, the interaction analysis had limited sensitivity for small effects; therefore, nonsignificant interaction effects were not interpreted as evidence of equivalent treatment responses among accessions.

To assess the potential influence of multimapping reads on candidate-gene expression estimates, an additional sensitivity analysis was performed using the same aligned BAM files and gene annotation used for the primary RNA-seq analysis. Gene-level read counts were regenerated with Rsubread::featureCounts in paired-end mode, but with multimapping reads excluded (countMultiMappingReads = FALSE). All other counting parameters were kept consistent with the primary analysis. For the nine candidate genes examined in Table 2, the total uniquely assigned read count was compared with the corresponding count from the original analysis, and the percentage of retained reads was calculated as (unique-only count/original count) × 100. This analysis was used to evaluate the robustness of candidate-gene expression estimates to multimapping and was not used to replace the primary transcriptome-wide count matrix.

**Table 1.**
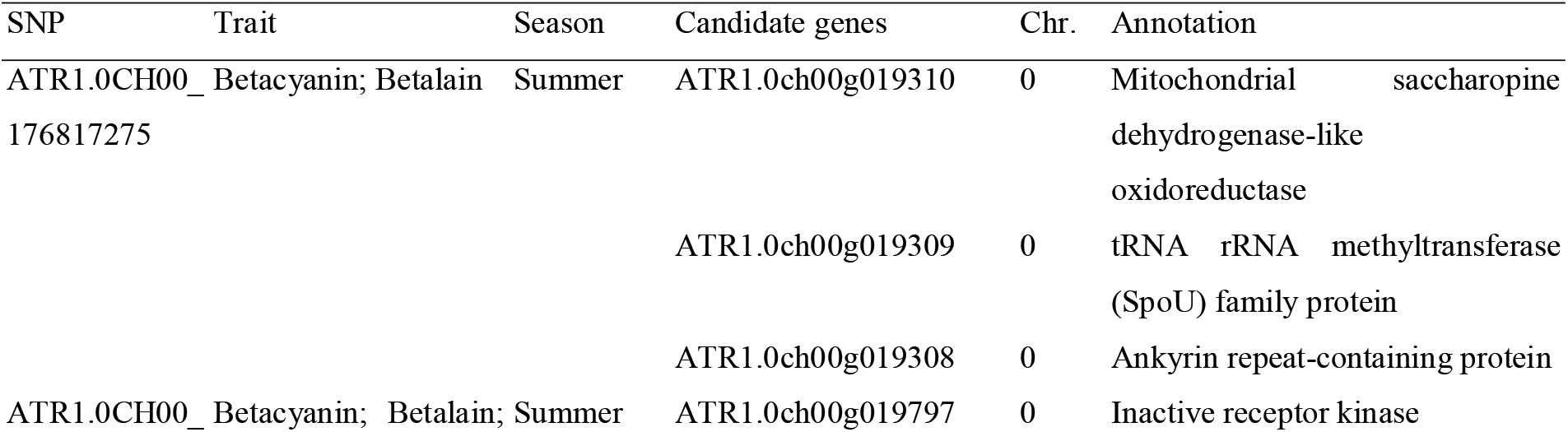
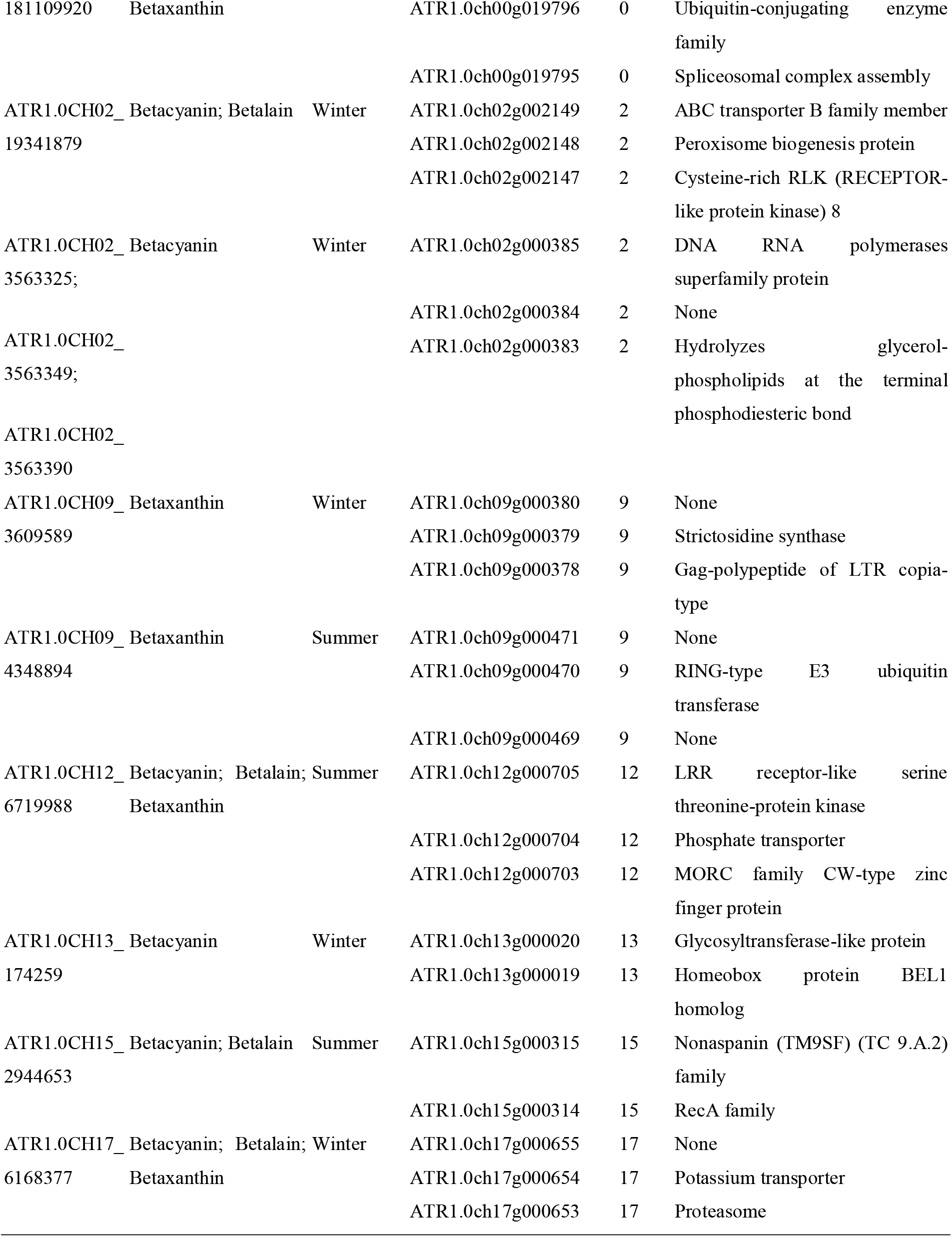
Twelve significant SNPs detected consistently by both FarmCPU and Blink models.

**Table 2.** Differential expression of the candidate genes under control and heat stress conditions.

| Gene | DE Homologs/ | Bayam |  | Am087 |  | AM127 |  |
| --- | --- | --- | --- | --- | --- | --- | --- |
|  | GWAS | log2FC | P value | log2FC | P value | log2FC | P value |
| ATR1.0ch02g002657 | 5GT | -2.27 | 1.00 | -4.91 | 0.00 | -1.99 | 1.00 |
| ATR1.0ch04g000541 | 5GT | -1.83 | 1.00 | -5.95 | 0.00 | -2.47 | 0.41 |
| ATR1.0ch09g000742 | 5GT | -1.81 | 1.00 | -2.79 | 0.00 | -2.25 | 0.71 |
| ATR1.0ch00g000258 | cDOPA | 1.38 | 1.00 | 3.84 | 0.00 | 5.52 | 0.00 |
| ATR1.0ch05g001519 | cDOPA | 1.38 | 1.00 | 3.83 | 0.00 | 5.51 | 0.00 |
| ATR1.0ch00g027747 | CYP76AD1 | -0.46 | 1.00 | -2.43 | 0.03 | -0.19 | 1.00 |
| ATR1.0ch03g000565 | ATR1.0CH03_5131103 | -1.49 | 1.00 | -1.31 | 1.00 | -3.77 | 0.05 |
| ATR1.0ch06g001201 | ATR1.0CH06_11090793 | -1.46 | 1.00 | -1.65 | 1.00 | -4.77 | 0.00 |
| ATR1.0ch13g000020 | ATR1.0CH13_174259 | -2.06 | 1.00 | -1.20 | 1.00 | -3.06 | 0.01 |

Gene Ontology (GO) enrichment analysis was conducted using TBtools-II (Chen et al., 2023), applying a Benjamini-Hochberg adjusted p-value threshold of < 0.05. The cis-elements analysis was conducted using the 5000 bp upstream sequences of candidate genes that exhibited differential expression under control and heat stress conditions (Table 2). Upstream (5**□**kb) motif scans identify candidate binding sites only; motif presence does not imply binding or regulation *in vivo*. The analysis was conducted using the New PLACE database, a repository of plant cis-acting regulatory DNA elements (https://www.dna.affrc.go.jp/PLACE/?action=newplace) (Higo et al. 1999).

### Quantitative real-time polymerase chain reaction analysis

Amaranthus accessions Bayam, Am087, and Am127 were grown under the same cultivation conditions as those used for RNA-seq. Leaves from one-month-old seedlings were harvested and immediately frozen in liquid nitrogen. Total RNA was extracted using the RNeasy Plant Mini Kit (Qiagen), following the manufacturer’s instructions. RNA quality and concentration were assessed by spectrophotometry (NanoDrop, Thermo Fisher Scientific) and confirmed by agarose gel electrophoresis. First-strand cDNA was synthesized from 0.2 µg of total RNA using the ReverTra Ace® qPCR RT Master Mix with gDNA Remover (TOYOBO), following standard protocols. Nine genes putatively involved in specialized metabolism or stress responses were selected based on previous transcriptome annotations and functional predictions (Table 2). These included three *5GT*-like genes, two *cDOPA*-like genes, one *CYP* gene, and three other genes. Gene identifiers were obtained from the Amaranthus genome database (e.g., *5G*: *ATR1.0ch02g002657*, *cDOPA*: *ATR1.0ch00g000258*, etc.). Gene-specific primers were designed using the GENETYX software (GENETYX Corp., Tokyo, Japan) and synthesized by Hokkaido System Science Co., Ltd. Details regarding the primers are provided in Supplementary Table 4. qRT-PCR was performed using a THUNDERBIRD^®^ SYBR™ qPCR Mix (TOYOBO) on a StepOnePlus Real-Time PCR System (Applied Biosystems). The thermal cycling conditions were as follows: initial denaturation at 95°C for 10 min, followed by 40 cycles of 95°C for 15 s and 60°C for 60 s. Primer specificity was verified by melting curve analysis. All samples were analyzed in triplicate, with three biological replicates per sample. The *actin-7* gene was used as an internal reference to calculate the relative unigene expression levels (Xiao et al., 2021). Expression data were analyzed using Microsoft Excel. Mean values and standard deviations were calculated, and significance of expression differences between accessions was assessed using one-way ANOVA followed by Tukey’s HSD test (p < 0.05).

## Results

### Betalain Absorbance Index Across Seasons

UV-Vis spectrophotometry was applied to a large panel, revealing overall variation in the pigment abundance across different seasons. During the winter, the betalain absorbance index (BAI = A538+A480; arbitrary absorbance units) ranged from 66.95 to 305.10, while in summer it ranged from 14.51 to 51.36, highlighting a general reduction in pigment□related absorbance under the warmer greenhouse season. The Pearson correlation coefficient between betalain-related absorbance indices in the two seasons was 0.29 (*p-value* = 0.006), suggesting a moderate but statistically significant correlation. Notably, red amaranths demonstrated higher betacyanin-related absorbance in winter compared to green amaranths, while both types showed the lowest betacyanin-related absorbance in summer (Supplementary Figure 1). In contrast, betaxanthin-related absorbance indices were primarily influenced by seasonal changes (Supplementary Figure 1), indicating that red and green amaranths may employ distinct regulatory mechanisms for betacyanin and betaxanthin synthesis in response to elevated temperatures.

### Reference Genome of *A. tricolor* cv. Bayam

The reference genome for *A. tricolor* cv. Bayam was constructed using a combination of 126.7 Gb high-quality Illumina short reads and 57.6 Gb PacBio Continuous Long Reads. Genome size estimation, based on *K*-mer distribution analysis, was approximately 596.4 Mb (Supplementary Figure 2); the assembly□to□estimate mismatch likely reflects repeat□rich regions and/or alternative haplotigs. We added this limitation explicitly and will improve anchoring with additional long□range data in future work.. PacBio sequencing yielded 57.6 Gb of reads with an N50 length of 38.1 kb, which were assembled into 2,151 contigs (Supplementary Table 5). A genetic map anchored 199 contigs to 17 chromosome-level scaffolds, spanning a total of 231.4 Mb, corresponding to 49.4% of the 468.43□Mb assembled span; 236.99□Mb remain unanchored (Supplementary Table 6). Repetitive sequences accounted for 52.9% of the genome, with LTR retroelements being the most dominant repeat type (8.9%), followed by simple repeats (3.9%) and DNA transposons (3.6%). In addition, a comprehensive sequence similarity search across the whole genome identified 5,599 pairwise homologous genes, forming 148 collinear blocks across 68 chromosome pairs (Supplementary Figure 3). Six pairs of these blocks were intra-chromosomal on chromosomes 1, 2, and 4, while the remaining 62 pairs were inter-chromosomal. Chromosome 2 exhibited 71 collinear blocks with all other chromosomes except for chromosome 16, whereas chromosome 14 shared only two homologous blocks with chromosome 2.

A total of 58,844 genes were annotated across the genome, including five genes encoding enzymes involved in betalain biosynthesis: *5GT*, *cDOPA*, *CYP76AD1*, *DODA*, and *TyDC*. These genes were blasted against the new reference genome to identify their homologous counterparts in *A. tricolor* (Supplementary Table 7). This search revealed 14, 18, 21, 2, and 5 homologous genes for *5GT*, *cDOPA*, *CYP76AD1*, *DODA*, and *TyDC*, respectively. Interestingly, *5GT* and *cDOPA* shared 12 homologous genes due to a 30% similarity in protein sequence (*E-value* = 10^-64). These homologous genes were assigned to either *5GT* or *cDOPA* based on the *E-value*, resulting in 12 and 8 *5GT* or *cDOPA*-like homologous genes, respectively.

### Marker-Trait Associations for Pigmentation-Related Absorbance Indices

In this core collection, a total of 40K SNPs were initially identified, with 2,912 SNPs retained following filtering based on minor allele frequency. Given the relatively limited marker density, one SNP per 200 Kb (Supplementary Figure 4), the detected associations are expected to span broad QTL intervals. Together with the observed LD-decay pattern (Supplementary Figure 5), these results should therefore be interpreted as tentative marker–trait associations and hypothesis-generating loci rather than fine-mapped QTLs. Principal Coordinate Analysis (PCA) revealed three genetic groups with several admixed accessions, indicating the need to account for population structure in the association analysis (Supplementary Figure 6). After the correction for population structure, the risk of false-positive associations was reduced (Supplementary Figure 7). GWAS identified 39 and 6 significant SNPs associated with betalain absorbance indices (A538, A480, and A538+A480) in the winter and summer seasons, respectively (Figure 3; Supplementary Figure 8; Supplementary Table 6). All the GWAS results of significant SNPs, including *p* values, adjusted *p* values, and effects were listed in Supplementary Table 8. Twelve marker-trait associations, seven in winter and five in summer, were consistently detected by both the FarmCPU and Blink models, providing cross-model support for these association signals. (Table 1). The SNP ATR1.0CH17_6168377 was identified in association with both betacyanin and betaxanthin during the winter season, while ATR1.0CH00_181109920 and ATR1.0CH12_6719988 were detected in association with both pigments during the summer season. These findings indicate that different marker–trait associations were detected across the two seasonal environments. Because genotype-by-environment interaction was not formally tested, these differences are interpreted as season-specific associations rather than evidence of significant G×E interaction. Putative positional candidates, comprising the SNP-containing gene when applicable and the immediately adjacent flanking genes, are listed in Supplementary Table 9. Notably, none of the 115 candidate genes identified have been previously implicated in betalain biosynthesis, indicating the need for further investigation into their potential roles.

**Figure 3.**
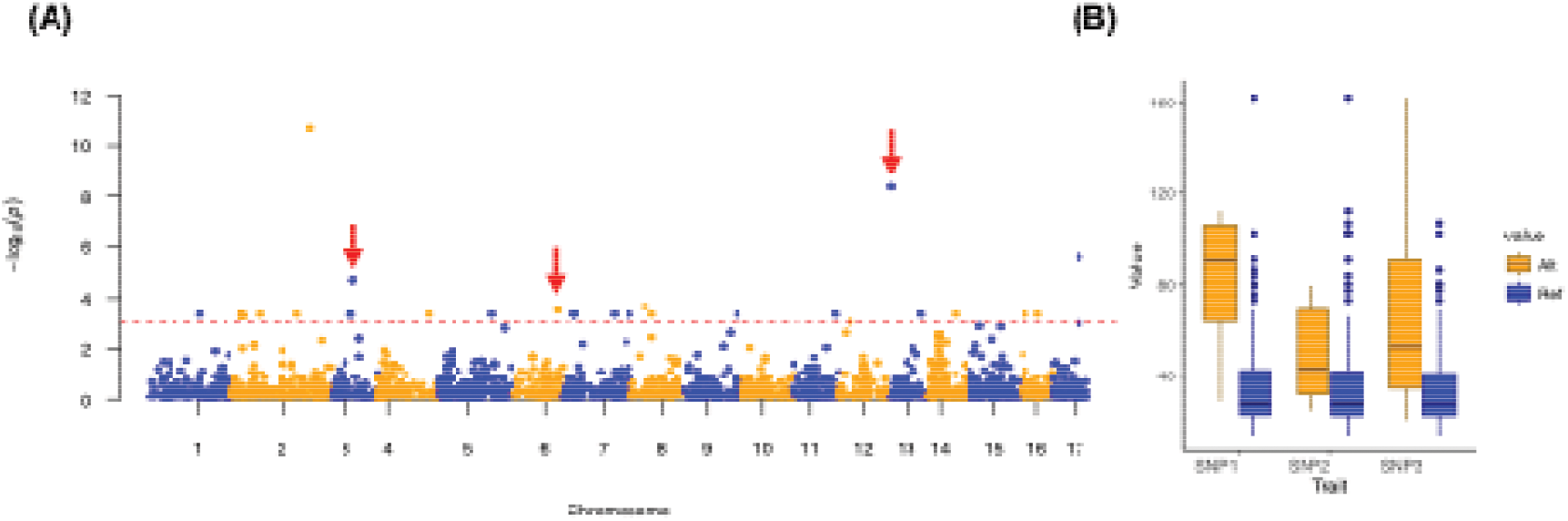
GWAS results of the betacyanin-dominanted absorbance. (A) Manhattan plot for betacyanin-dominated absorbance in the winter season using the FarmCPU model. The horizontal red dashed line indicates the significance cutoff (*p* < 0.001). Red arrows mark significant SNPs co-localized with DEGs. (B) Allelic effects of the SNPs co-localized with DEGs (SNP1: ATR1.0CH03_5131103; SNP2: ATR1.0CH06_11090793; SNP3: ATR1.0CH13_174259). Box plots illustrate phenotypic distributions by genotype class; the blue and orange boxes represent the reference and alternative allele homozygotes, respectively.

### Differential Gene Expression Under Heat Stress

To investigate the genetic mechanisms underlying betalain biosynthesis under varying temperature conditions, RNA-seq experiments were conducted on Am087 and Am127, two accessions that displayed relatively consistent performance in field trials and higher betalain-related absorbances across seasons. The cultivated line Bayam was used as a control. One month after sowing, plants subjected to heat stress exhibited typical stress phenotypes, including stem elongation, delayed panicle production, reduced pigmentation, leaf abscission, senescence, and curling (Figure 4). Pigmentation-related absorbance indices decreased in Am087 and Am127 but remained relatively stable in Bayam. (Supplementary Table 10). This was consistent with the similar changes of betalain-related absorbance in Am087 and Am127.

**Figure 4.**
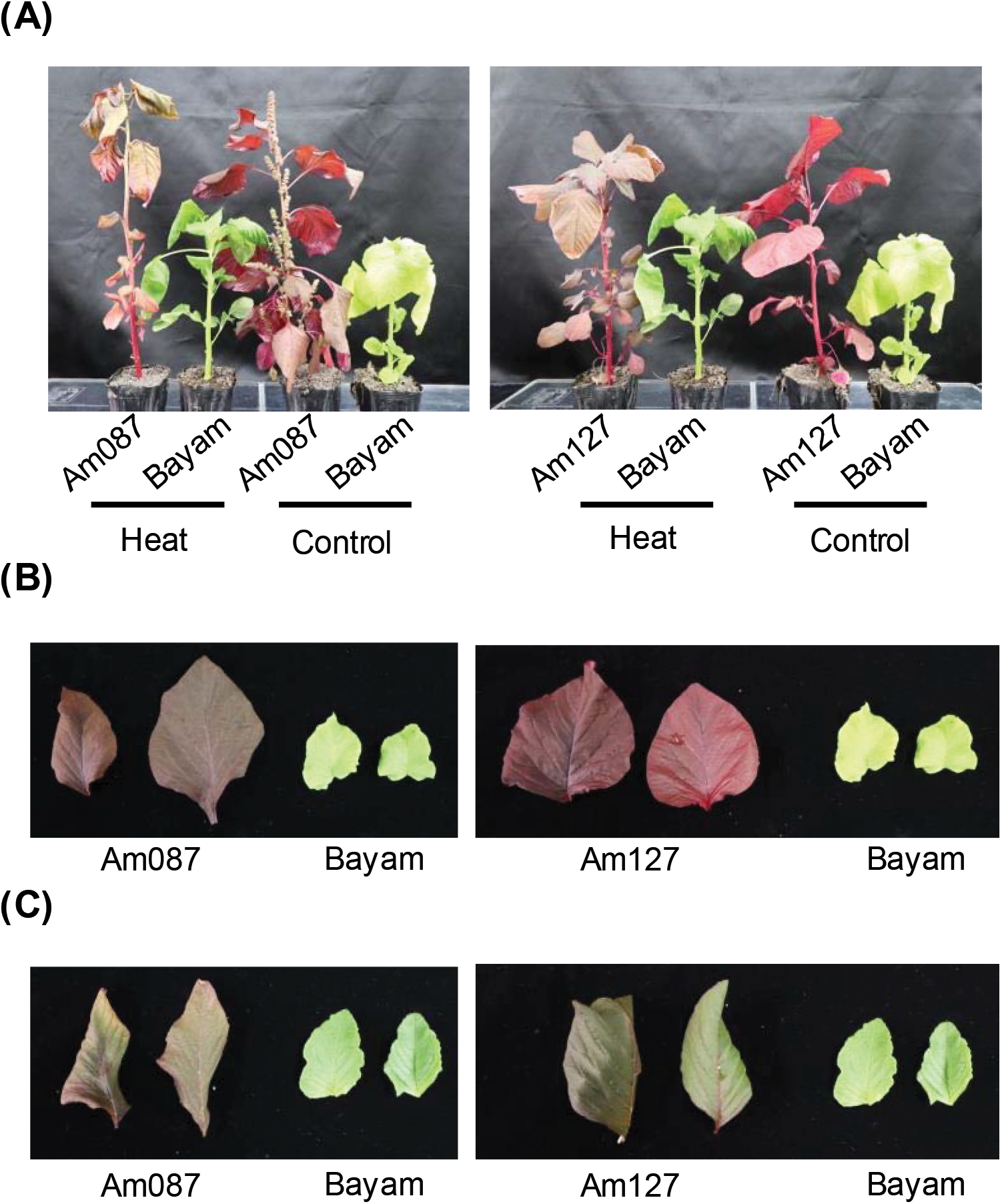
Morphological comparison of Bayam, Am087, and Am127 under control and heat stress conditions. (A) Whole-plant morphology of Bayam, Am087, and Am127 under control (25°C) and heat stress (35°C day/25°C night) conditions. (B) Leaf color variation under control conditions. (C) Leaf color variation under heat stress. Plants were grown under a 16 h light/8 h dark photoperiod for one month. Heat stress was applied for two weeks before sampling. Images were taken before betalain extraction.

Across the 18 libraries included in the final analysis, 97.53–98.50% of primary reads were mapped to the reference genome, and 96.59–97.81% of reads were properly paired. In addition, 81.35–85.77% of fragments were assigned to annotated genes, while 0.84–1.48% were classified as unmapped, 11.83–15.18% did not overlap annotated features, and 1.18–2.63% were assigned ambiguously. PCA of the variance-stabilized expression profiles showed that PC1 and PC2 explained 33.3% and 26.4% of the total variance, respectively (Supplementary Figure 9A). Sample-to-sample Pearson correlation analysis showed generally high correlations among biological replicates within the same genotype × treatment group (Supplementary Figure 9B). Dispersion diagnostics showed that most genes followed the fitted mean–dispersion relationship, with a median dispersion of 0.061 and an interquartile range of 0.025–0.186; the 95th percentile was 0.714, although a smaller subset of genes showed higher dispersion (Supplementary Figure 9C.) Bayam exhibited 3 DEGs (*adjusted p values*□<□0.05), whereas Am087 and Am127 showed 506 and 771 DEGs, respectively, (Supplementary Table 11). A total of 100 DEGs were shared between Am087 and Am127 (Figure 5A). Both lines exhibited enrichment in genes related to transmembrane transport activity (Supplementary Figure 10), consistent with the expected impact of heat stress on transmembrane stability.

**Figure 5.**
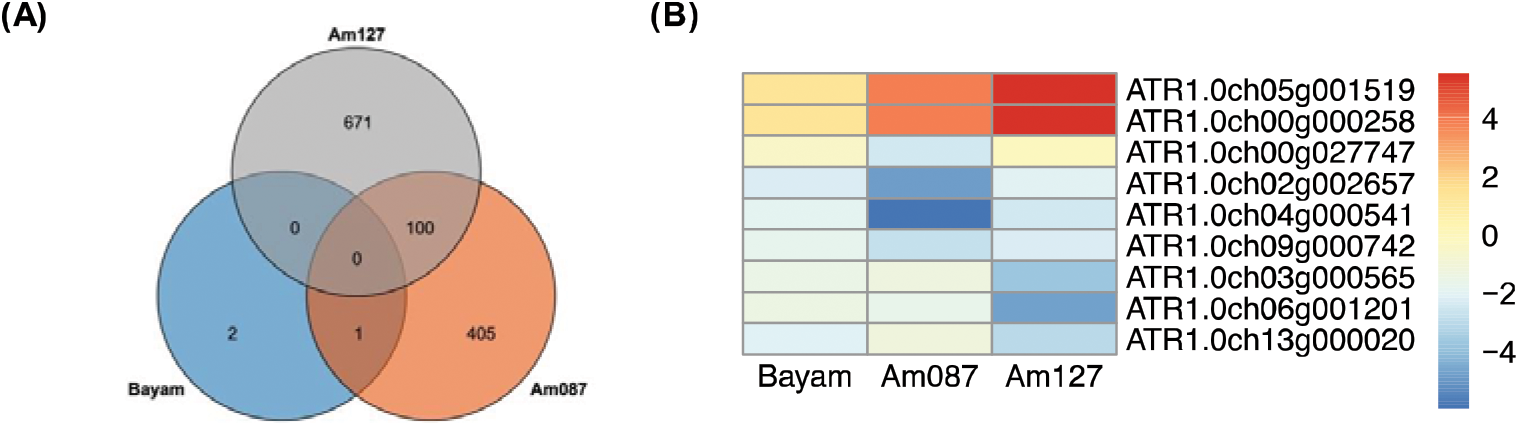
Transcriptomic profiles of the three lines. (A) Venn diagram showing differentially expressed genes (DEGs) among Bayam, Am087, and Am127 under heat stress. (B) Heat map showing the expression levels of candidate betalain biosynthesis genes, *ATR1.0ch05g001519* and *ATR1.0ch00g000258* are homologs of *cDOPA*, while *ATR1.0ch00g027747* is a homolog of *CYP76AD1*. Additionally, *ATR1.0ch02g002657*, *ATR1.0ch04g000541*, and *ATR1.0ch09g000742* are homologs of *5GT*. The genes *ATR1.0ch03g000565*, *ATR1.0ch06g001201*, and *ATR1.0ch13g000020* are DEGs associated with significant SNPs under optimal conditions. In the heat map, a redder color indicates higher expression levels under heat stress, while a bluer color indicates lower expression levels under control conditions.

To formally evaluate whether transcriptional responses differed among accessions, a genotype × treatment interaction analysis was performed. The interaction test identified 4,638 genes with significant genotype-dependent treatment responses (FDR < 0.05), while pairwise interaction contrasts identified 1,156 and 2,556 genes with differential treatment responses between Am087 and Bayam and between Am127 and Bayam, respectively. Because each genotype × treatment combination contained three biological replicates, the interaction analysis had limited sensitivity for small effects; nonsignificant interaction results were therefore interpreted cautiously and were not considered evidence of equivalent transcriptional responses among accessions.Six homologous genes involved in betalain biosynthesis—two *cDOPA*-like homologs (*ATR1.0ch05g001519*, *ATR1.0ch00g000258*), one *CYP76AD1* homolog (*ATR1.0ch00g027747*), and three *5GT* homologs (*ATR1.0ch02g002657*, *ATR1.0ch04g000541*, *ATR1.0ch09g000742*)—were differentially expressed in Am087 and Am127 during summer. In Am087, *CYP76AD1* was significantly downregulated, while the *5GT* homologs were also downregulated, despite the upregulation of *cDOPA*-like homologs (Figure 5B, Table 2). Am127 exhibited similar trends, although without statistical significance (Table 2). These results indicate an association between reduced pigmentation-related absorbance and decreased expression of *CYP76AD1* and *5GT* homologs under elevated-temperature conditions, with the upregulation of *cDOPA* unable to compensate for this decrease.

To evaluate the expression profiles of candidate genes potentially involved in metabolite biosynthesis and physiological traits, we analyzed nine genes, including six putative homologs involved in betalain biosynthesis (three *5GT*: *ATR1.0ch02g002657*, *ATR1.0ch04g000541*, *ATR1.0ch09g000742*, two *cDOPA*: *ATR1.0ch00g000258*, *ATR1.0ch05g001519*, one *CYP*: *ATR1.0ch00g027747*) and three additional DEGs (*ATR1.0ch03g000565*, *ATR1.0ch06g001201*, and *ATR1.0ch13g000020)* identified through GWAS integration in three Amaranthus accessions: Bayam, Am087, and Am127 (Table 2, Supplementary Figure 11). Distinct expression patterns were observed among the accessions for several genes. *ATR1.0ch02g002657* encoding *5GT* showed markedly higher expression in Bayam compared to Am127, suggesting accession-specific regulation. Similarly, *ATR1.0ch00g000258* encoding *cDOPA* exhibited a significant increase in transcript levels in Am087, whereas ATR1.0ch00g027747 encoding *CYP* was expressed at lower levels in Bayam relative to both Am087 and Am127. The remaining genes also demonstrated variable expression across the accessions, although the magnitude of difference varied. These data suggest that the expression of these genes is differentially regulated depending on the genetic background of each accession and is broadly consistent with the RNA-seq results.

Because *ATR1.0ch00g000258* and *ATR1.0ch05g001519* showed nearly identical RNA-seq expression profiles and share extremely high sequence similarity, we evaluated the effect of multimapping reads on their quantification. Recounting the RNA-seq data after excluding multimapping reads retained 98.3–100% of the original counts for the other seven candidate genes examined, whereas only 1.18% and 0.06% of the original counts were retained for *ATR1.0ch00g000258* and *ATR1.0ch05g001519*, respectively (Supplementary Table 12). These results indicate that the original short-read RNA-seq data cannot reliably resolve expression of the two *cDOPA*-like loci independently. Therefore, their nearly identical expression profiles are interpreted as a *cDOPA*-like transcript signal rather than as independent locus-specific responses.

Considering that our SNP density is approximately one marker per 200 Kb, substantially lower than the average gene density of ∼1 per 10 Kb, we integrated DEGs to refine GWAS candidate genes. By anchoring statistical associations to heat-responsive DEGs, the likelihood of false positives was reduced, improving biological relevance. This approach is based on the rationale that a gene associated with betalain biosynthesis in only one season, and concurrently identified as a DEG, is likely involved in the regulation of betalain synthesis during that specific season. Three significant SNPs identified in the winter betacyanin GWAS were associated with three DEGs (*ATR1.0ch03g000565*, *ATR1.0ch06g001201*, *ATR1.0ch13g000020*), which were significantly downregulated in Am127 under heat stress (Figure 5B, Supplementary Table 11).

These three genes represent promising candidate genes associated with variation in pigmentation-related traits under elevated-temperature conditions. In addition, 5000 bp upstream sequences of candidate genes showing differential expression under control and heat stress conditions were hypothesized to contain promoter and cis-element sequences (Table 2). Cis-element analysis using the New PLACE database identified the presence of regulatory elements in multiple candidate genes, including *ATR1.0ch02g002657*, *ATR1.0ch04g000541*, *ATR1.0ch09g000742*, *ATR1.0ch00g000258*, *ATR1.0ch05g001519*, *ATR1.0ch00g027747*, *ATR1.0ch03g000565*, *ATR1.0ch06g001201*, and *ATR1.0ch13g000020*. These genes were associated with 74, 71, 57, 122, 178, 109, 159, 72, and 71 signal sequences, respectively, related to MYB and WRKY regulation, respectively (Supplementary Table 13).

## Discussion

### Putative Candidate Genes Prioritized from GWAS Signals

We integrated the tentative GWAS signals with differential-expression data to prioritize biologically plausible candidate genes. Given the limited marker density, these genes should be regarded as putative candidates associated with hypothesis-generating loci rather than fine-mapped causal genes. An significant SNP, ATR1.0CH14_4712147, is located within *ATR1.0ch14g000595*, which encodes a DEAD-box ATP-dependent RNA helicase. DEAD-box RNA helicases are crucial for plant development, influencing both vegetative and reproductive growth, as well as mediating responses to abiotic and biotic stress (Li et al., 2023). For example, *OsBIRH1* in rice encodes a DEAD-box RNA helicase that is vital for tolerance to both biotic and abiotic stresses (Li et al., 2008). Similarly, the *Thermotolerant Growth Required 1* in Chinese cabbage has been associated with heat stress tolerance (Yarra and Xue, 2020). In Arabidopsis, T-DNA insertion mutants of DEAD-box RNA helicases show upregulation of *MYB* genes under abiotic stress. *MYB* transcription factors are known to regulate anthocyanin biosynthesis (Petroni and Tonelli, 2011; Xue et al., 2025). The *MYB* gene, which directly regulates both *CYP76AD1* and *DODA* in *B. vulgaris* (Hatlestad et al., 2015), is located on Chromosome 2 and is linked to the *CYP76AD1* locus (Goldman and Austin, 2000). In pitaya, *HuMYB132* has been demonstrated to promote betacyanin biosynthesis by binding to *HuCYP76AD1–1* and *HuDODA1* (Xie et al., 2023). In this study, seven *MYB* family genes were significantly differentially expressed in Am127, with six of them downregulated during the summer season. Although *ATR1.0ch14g000595* was not identified as a DEG, genetic variation at this locus may potentially alter its structure, leading to the downregulation of *MYB* genes. Furthermore, earlier research has indicated that *WRKY* genes regulate pigment biosynthesis by interacting with *MYB* genes (Li et al., 2020; Xie et al., 2023). However, in this study, we identified three *WRKY* genes as DEGs, all of which were upregulated under heat stress conditions. This finding contrasts with previous results, suggesting a complex regulatory mechanism governing betalain biosynthesis under abiotic stress. In our dataset, seven *MYB* family genes were differentially expressed in Am127, with six downregulated under heat stress. Additionally, three *WRKY* genes were upregulated. Cis-element analysis of candidate betalain genes revealed multiple *MYB* and *WRKY* binding motifs (Supplementary Table 13). These findings provide species-specific evidence suggesting the potential involvement of *MYB* and *WRKY* transcription factors in pigmentation-related regulation in *A. tricolor*. However, functional binding and regulatory activity remain to be demonstrated. Further validation using chromatin immunoprecipitation (ChIP) and reporter assays is necessary to confirm their regulatory roles. Notably, the current resolution (∼1 marker per 200 kb) may be insufficient to capture fine-scale genetic loci. Therefore, future studies should employ higher-resolution sequencing or targeted SNP marker development.

### Heat Stress Reduces betalain-related absorbance in *Amaranthus tricolor*

This study found lower pigmentation-related absorbance in the summer seasonal environment than in winter, consistent with previous findings in amaranth under elevated-temperature conditions (Reyes-Rosales et al., 2023). Because the seasonal trials differed in multiple environmental factors, these differences cannot be attributed specifically to temperature. In the controlled-environment experiment, Bayam showed relatively smaller changes in pigmentation-related absorbance than Am087 and Am127 under the elevated-temperature regime. However, because physiological and growth-related indicators of heat tolerance were not evaluated, this response should be interpreted specifically as relative stability of the pigmentation-related phenotype rather than evidence of whole-plant thermotolerance. The candidate genes identified here, including *CYP76AD1* and *5GT* homologs, therefore provide targets for further functional investigation of pigmentation responses to elevated-temperature conditions.

While UV-V is spectrophotometry provides a practical assessment of pigment abundance, absorbance values should not be interpreted as direct measures of betalain-related absorbance and must be viewed with appropriate caution; future studies incorporating component-specific analytical approaches such as HPLC or multi-wavelength spectral deconvolution will enable more precise resolution of betacyanin and betaxanthin composition. In this study, we treat all locus□level results as hypothesis□generating. In addition, we evaluated the betalain absorbance index across a large panel under greenhouse conditions by leveraging natural temperature differences between summer and winter. Our seasonal comparison reflects environmental co□variation; accordingly, we do not ascribe pigment changes solely to heat. We now interpret these results as environment□associated changes rather than definitive heat□stress effects. It should be noted that additional environmental factors, such as daytime temperature maxima, VPD, irradiance, and photoperiod, may also influence betalain biosynthesis and accumulation. Similarly, in the growth-chamber assays, the 35□°C temperature regime may have been accompanied by elevated VPD due to uncontrolled relative humidity. Therefore, future studies should incorporate increased replication or more tightly controlled experimental designs to better disentangle the effects of individual environmental variables. From an applied perspective, heat stress in agricultural systems commonly occurs in combination with other abiotic stresses. Accordingly, our approach reflects realistic heat-stress scenarios encountered under conventional agricultural practices, thereby providing results with greater ecological and practical relevance.

### The Disruption of Betalain Biosynthesis Under Heat Stress Conditions

Previous studies have established that *CYP76AD1* is crucial for the initial step of betalain biosynthesis, which proceeds either through *DODA* and *5GT* or *CYP76AD1* and *cDOPA* to produce betanin, and via *DODA* to synthesize betaxanthin (Chang et al., 2021). In the current study, both RNA-Seq and qRT-PCR analyses demonstrated that *CYP76AD1* homologs (ATR1.0ch00g027747) were consistently downregulated under elevated temperature, particularly in Am087, where transcript levels were significantly lower than in Bayam and Am127; these transcript□level associations do not establish enzymatic or metabolic causality. These results indicate an association between reduced pigmentation-related absorbance and decreased expression of *CYP76AD1* and *5GT* homologs under elevated-temperature conditions. Similar downregulation of *CYP76AD1* under high temperatures has been documented in *Suaeda salsa* (Li et al., 2023), suggesting that this may be a common mechanism for the degradation of betalain biosynthesis across species. The *5GT* homologs (*ATR1.0ch02g002657*, *ATR1.0ch04g000541*, *ATR1.0ch09g000742*), which contribute to the final glycosylation step in betacyanin biosynthesis, were also downregulated under heat stress, especially in Am087 and Am127, as validated by qRT-PCR. In contrast, *cDOPA*-like homologs (*ATR1.0ch00g000258*, *ATR1.0ch05g001519*) were upregulated, most notably in Am087. However, this increase was insufficient to offset the reduction in betacyanin-dominated absorbance observed in Am087 under stress conditions. Interestingly, RNA-Seq and qRT-PCR analyses revealed that Bayam maintained stable expression of *CYP76AD1* and *5GT* under heat stress, in contrast to Am087 and Am127, which showed significant downregulation. This transcriptional stability may contribute to the maintenance of pigmentation-related absorbance in Bayam and suggests a resilient regulatory network. These accession-specific differences may reflect variation in promoter architecture, transcription factor activity, or epigenetic regulation, positioning Bayam as a promising donor for breeding heat-tolerant cultivars. In contrast, Am087 exhibited the most pronounced suppression of both genes, aligning with its marked reduction in pigmentation-related absorbance. This differential gene response between accessions indicates the involvement of accession-specific regulatory mechanisms, potentially driven by upstream transcription factors or signaling components. The expression of *DODA*-another key enzyme in the betaxanthin branch-did not differ significantly between control and heat stress conditions, suggesting that *CYP76AD1* may contribute to the observed reduction in pigmentation-related absorbance under elevated-temperature conditions. Collectively, these findings highlight *CYP76AD1*, *5GT*, and *cDOPA*-like homologs as candidate genes associated with pigmentation-related responses under elevated-temperature conditions. Importantly, the disproportionate decline in betacyanin-dominated absorbance observed in Am087 compared to Am127 suggests that additional transcriptional repressors or chromatin-level modifications may contribute to the accession-specific sensitivity of betalain biosynthesis to heat. While transcriptomic and qRT-PCR analyses provide insights into gene expression changes, future studies should integrate proteomic and metabolomic approaches to confirm functional changes at the protein and metabolite levels. Future studies will explore stress mitigation strategies to counteract betalain reduction under heat stress. These strategies may include exogenous application of signaling molecules, such as salicylic acid, or genetic modifications to enhance *CYP76AD1* and *5GT* expression. This study has several limitations. First, pigmentation traits were evaluated using absorbance-based indices rather than chromatographic quantification. Second, physiological indicators of heat stress were not measured. Third, the marker density of the GWAS dataset was relatively low. Finally, the candidate genes identified here require functional validation through genetic and biochemical approaches.

## Conclusions

We generated a reference genome for the cultivar Bayam and integrated genome-wide association studies (GWAS) with transcriptomic analyses to identify candidate loci associated with pigmentation-related traits in *Amaranthus tricolor*. GWAS identified multiple genomic regions associated with variation in pigment-related absorbance indices across seasonal environments. Integration of GWAS and differential expression analyses highlighted several candidate genes, including *ATR1.0ch03g000565*, *CYP76AD1*, and *5GT* homologs, as promising targets for future investigation. Reduced pigmentation-related absorbance under elevated-temperature conditions was associated with decreased expression of *CYP76AD1* and *5GT* homologs, whereas *cDOPA*-like homologs exhibited increased expression. Given the absorbance-based nature of the pigmentation measurements and the limited resolution of the GWAS dataset, these findings should be considered hypothesis-generating rather than definitive evidence of causality. Collectively, the genomic resources generated in this study provide a foundation for future functional analyses and marker-assisted breeding of climate-resilient amaranth cultivars.

## Data availability

Raw sequence reads were deposited in the Sequence Read Archive (SRA) database of the DNA Data Bank of Japan (DDBJ) under the accession number DRA014927, DRA014962, and DRA014963. Assembled sequences are available at DDBJ (accession numbers BSCK01000001 - BSCK01001969) and Plant GARDEN (https://plantgarden.jp).

## Acknowledgments

This work was supported by JSPS KAKENHI (Grant Numbers 19KK0151, 22H05172, and 22H05181) and the Kazusa DNA Research Institute Foundation. We would like to thank National Core Facility for Biopharmaceuticals (NCFB, NSTC 114-2740-B-492-001) and National Center for High-performance Computing (NCHC) of National Institutes of Applied Research (NIAR) of Taiwan for providing computational and storage resources. We thank the staff in Kazusa DNA Research Institute for their technical support. We also thank the long-term strategic donors to the World Vegetable Center: Taiwan, the United States, Australia, the United Kingdom, Germany, Thailand, South Korea, the Philippines, and Japan.

## Author contributions statement

Y.-P. Lin designed and performed research, analyzed data, and wrote the paper. T. Ohokubo performed research. A. Nashiki performed research and analyzed data. Y. Yoshioka performed research and analyzed data. Y.-Y. Zhang performed research. H.-M. Ting performed research. Sachiko Isobe designed and performed research. Kenta Shirasawa designed and performed research and wrote the paper. Ken Hoshikawa designed and performed research, analyzed data, and wrote the paper.

## Additional Information

Competing financial interests.

The authors declare that they have no competing interests.

**Abbreviations**
CYP76AD1: cytochrome P450 76AD1
L-DOPA: L-3,4-dihydroxyphenylalanine
DODA: L-dihydroxyphenylalanine 4,5-dioxygenase
cDOPA5GT: cyclo-DOPA 5-O-glucosyltransferase
B5GT: betanidin-5-O-glucosyltransferase
TyDC: tyrosine decarboxylase

## Legends of supplementary data

**Supplementary Figure 1 Comparisons of betalain between seasons and leaf colors.** The error bars indicate the standard errors. The letters upon the bars refer to the statistical groups resulting from the Tukey’s HSD test.

**Supplementary Figure 2 The genome size of Bayam.** *K*-mer analysis indicates the genome size of Bayam is about 596.4 Mb

**Supplementary Figure 3 Genome duplication in the *A. tricolor* cv. Bayam. Similar genome segments are connected with lines.**

**Supplementary Figure 4 Visualization of SNP Distribution.** This figure illustrates the distribution of single nucleotide polymorphisms (SNPs) across 17 chromosomes. The horizontal axis represents chromosomal positions in megabases (Mb), while the vertical axis lists chromosome numbers. Color intensity indicates SNP density, ranging from green (low density) to red (high density), as shown in the legend (0–900). Regions in red correspond to high SNP concentration, whereas green and yellow areas indicate lower density. This distribution pattern provides valuable insights for genetic diversity analysis, quantitative trait locus (QTL) mapping, and genomic selection.

**Supplementary Figure 5 Genome-wide linkage disequilibrium (LD) decay pattern.** The plot illustrates the decline of LD with increasing physical distance across the genome. The x-axis represents physical distance in megabases (Mb), and the y-axis shows the pairwise LD measure (R^2) between SNPs. Gray dots indicate individual SNP pairs, while the purple line represents the sliding window average and the blue line shows the LOESS-smoothed LD decay curve. LD decreases rapidly within short distances and stabilizes beyond several hundred kilobases, reflecting the extent of recombination and genetic diversity in the population.

**Supplementary Figure 6 Principal Coordinate Analysis of the core collection.**Colors indicate the leaf color of each accession.

**Supplementary Figure 7 Q-Q plots of betaxanthin-dominated (A480), betacyanin-dominant (A538), and betalain absorbance index (A480 + A538).** A) and B) refer to the Q-Q plots of the FarmCPU and the Blink models, respectively.

**Supplementary Figure 8 Manhattan plots of betaxanthin-dominated (A480), betacyanin-dominant (A538), and betalain absorbance index (A480 + A538).** A) The Manhattan plots of the FarmCPU model. B) The Manhattan plots of the Blink model. The horizontal red dashed line indicates the significance cutoff (*p value*□<□0.001).

**Supplementary Figure 9 RNA-seq sample-level and DESeq2 model diagnostics.** (A) Principal component analysis of the 18 RNA-seq libraries based on variance-stabilized expression values of the 500 most variable genes. Colors indicate accession and symbols indicate treatment. PC1 and PC2 explain 33.3% and 26.4% of the total variance, respectively. (B) Sample-to-sample Pearson correlation heatmap calculated from variance-stabilized expression values across the 18 libraries. (C) DESeq2 dispersion diagnostic showing gene-wise dispersion estimates and the fitted mean–dispersion relationship.

**Supplementary Figure 10 GO enrichment analyses.** A) and B) are the top 30 enriched GO in Am087 and Am127, respectively. C) is the enriched GO shared by Am087 and Am127.

**Supplementary Figure 11 Relative expression patterns of candidate genes in different Amaranthus accessions under control and heat stress conditions.** (A) 5GT: *ATR1.0ch02g002657*, (B) 5GT: *ATR1.0ch04g000541*, (C) 5GT: *ATR1.0ch09g000742*, (D) cDOPA: *ATR1.0ch00g000258*, (E) cDOPA: *ATR1.0ch05g001519*, (F) CYP: *ATR1.0ch00g027747*, (G) *ATR1.0ch03g000565*, (H) *ATR1.0ch06g001201*, (I) *ATR1.0ch13g000020*. Relative expression levels were measured by qRT-PCR in leaves of Amaranthus accessions: Bayam, Am087, and Am127. The heat stress condition was as described in Materials and Methods. Gene expression was compared across three Amaranthus accessions: Bayam, Am087, and Am127. The bars represent the mean values (n = 3) and intervals over the bars indicate the standard deviation of the means. * Significant change (*p value* < 0.05) relative to the control.

**Supplementary Table 1 Phenotypes of the core collection under the two seasons.**

**Supplementary Table 2 Summary of ddRAD-seq reads and coverage per sample.**

**Supplementary Table 3 Summary of RNA-seq reads and coverage under control and heat stress.**

**Supplementary Table 4 qRT-PCR primer sets for the nine candidate genes in *Amaranthus*.**

**Supplementary Table 5 Statistics of the genome assembly of *A. tricolor* cv. Bayam.**

**Supplementary Table 6 Statistical assembly summary of each chromosome.**

**Supplementary Table 7 The homologous genes of the five enzymes in the *A. tricolor* genome.**

**Supplementary Table 8 Significant SNPs of the betalain-related absorbance indices in the two seasons detected by FarmCPU and Blink models in GAPIT.** Columns are: Trait (A538 = betacyanin□dominated absorbance; A480 = betaxanthin□dominated absorbance; BAI = A538 + A480, arbitrary absorbance units), Season, Model (BLINK/FarmCPU), SNP ID, Chromosome, Position (bp), P.value (raw), MAF (minor allele frequency), H.B.P.Value (Benjamini–Hochberg FDR□adjusted P), and Effect (additive effect estimated by the model; positive values indicate higher absorbance for the alternative allele). Unless otherwise noted, associations are interpreted as hypothesis□generating given marker density and LD. Sample size is n□=□95 accessions evaluated per season where phenotypes were available.

Notes: Traits are UV–Vis absorbance indices (relative, non□specific) rather than chromatographically quantified metabolites; see Materials and Methods for details of trait definitions and GWAS settings (models, kinship/PCs, multiple□testing control).

**Supplementary Table 9 Candidate genes directly targeted by or adjacent to the significant SNPs.**

**Supplementary Table 10 Betacyanin and betaxanthin of *A. tricolor* cv. Bayam, Am087, and Am127 under the control and heat conditions.**

**Supplementary Table 11 DEGs in Bayam, Am087, and Am127 and their annotations.**

**Supplementary Table 12 Sensitivity analysis of candidate-gene read counts after exclusion of multimapping reads.**

**Supplementary Table 13 Cis-elements related to *MYB* and *WRKY* on the candidate genes.**

## References

1. Albano C, Negro C, Tommasi N, Gerardi C, Mita G, Miceli A, De Bellis L, Blando F. 2015. Betalains, phenols and antioxidant capacity in cactus pear [*Opuntia ficus-indica* (L.) Mill.] fruits from Apulia (South Italy) genotypes. Antioxidants. 4(2): 269–280.

2. Bolger AM, Lohse M, Usadel B. 2014. Trimmomatic: a flexible trimmer for Illumina sequence data. Bioinformatics. 30(15): 2114–2120.

3. Bradbury PJ, Zhang Z, Kroon DE, Casstevens TM, Ramdoss Y, Buckler ES. 2007. TASSEL: software for association mapping of complex traits in diverse samples. Bioinformatics. 23(19):2633–2635.

4. Cantalapiedra CP, Hernández-Plaza A, Letunic I, Bork P, Huerta-Cepas J. 2021. eggNOG-mapper v2: functional annotation, orthology assignments, and domain prediction at the metagenomic scale. Molecular Biology and Evolution. 38(12): 5825–5829.

5. Chang YC, Chiu YC, Tsao NW, Chou YL, Tan CM, Chiang YH, Liao PC, Lee YC, Hsieh LC, Wang SY et al. 2021. Elucidation of the core betalain biosynthesis pathway in *Amaranthus tricolor*. Scientific Reports. 11(1): 6086.

6. Chen C, Wu Y, Li J, Wang X, Zeng Z, Xu J, Liu Y, Feng J, Chen H, He Y, Xia R. 2023. TBtools-II: A “one for all, all for one” bioinformatics platform for biological big-data mining. Mol. Plant. 16: 1733–1742.

7. Cox MP, Peterson DA, Biggs PJ. 2010. SolexaQA: at-a-glance quality assessment of Illumina second-generation sequencing data. BMC Bioinformatics. 11: 485.

8. FAO, IFAD, UNICEF, WFP, WHO. 2018. Africa regional overview of food security and nutrition: addressing the threat from climate variability and extremes for food security and nutrition. Addis Ababa. © UN. ECA. https://hdl.handle.net/10855/43032”

9. Galanie S, Thodey K, Trenchard IJ, Filsinger Interrante MF, Smolke CD. 2015. Complete biosynthesis of opioids in yeast. Science. 349(6252): 1095–1100.

10. Gremme G, Brendel V, Sparks ME, Kurtz S. 2005. Engineering a software tool for gene structure prediction in higher organisms. Information and Software Technology. 47(15): 965–978.

11. Goldman IL, Austin D. 2000. Linkage among the R, Y and BI loci in table beet. Theoretical and Applied Genetics. 100: 337–343.

12. Haas BJ, Salzberg SL, Zhu W, Pertea M, Allen JE, Orvis J, White O, Buell CR, Wortman JR. 2008. Automated eukaryotic gene structure annotation using EVidenceModeler and the Program to Assemble Spliced Alignments. Genome Biology, 9(1): R7.

13. Haas B, Papanicolaou A, Yassour M. 2016. TransDecoder. https://github.com/TransDecoder/TransDecoder.

14. Hatlestad GJ, Akhavan NA, Sunnadeniya RM, Elam L, Cargile S, Hembd A, Gonzalez A, McGrath JM, Lloyd AM. 2015. The beet *Y* locus encodes an anthocyanin MYB-like protein that activates the betalain red pigment pathway. Nature Genetics. 47(1): 92–96.

15. Higo K, Ugawa Y, Iwamoto M, Korenaga T. 1999. Plant cis-acting regulatory DNA elements (PLACE) database: Nucleic Acids Research. 27 (1): 297–300

16. Hoshikawa K, Lin Y-P, Schafleitner R, Shirasawa K, Isobe S, Nguyen DC, Ohsawa R, Yoshioka Y. 2023. Genetic diversity analysis and core collection construction for *Amaranthus tricolor* germplasm based on genome-wide single-nucleotide polymorphisms. Scientia Horticulturae. 307: 111428.

17. Kim D, Paggi JM, Park C, Bennett C, Salzberg SL. 2019. Graph-based genome alignment and genotyping with HISAT2 and HISAT-genotype. Nature Biotechnology. 37(8): 907–915.

18. Langmead B, Salzberg SL. 2012. Fast gapped-read alignment with Bowtie 2. Nature Methods. 9(4): 357–359.

19. Li C, Wu J, Hu K, Wei S, Sun H, Hu L, Han Z, Yao G, Zhang H. 2020. *PyWRKY26* and *PybHLH3* cotargeted the *PyMYB114* promoter to regulate anthocyanin biosynthesis and transport in red-skinned pears. Horticulture Research, 7: 37

20. Li D, Liu H, Zhang H, Wang X, Song F. 2008. OsBIRH1, a DEAD-box RNA helicase with functions in modulating defence responses against pathogen infection and oxidative stress. Journal of Experimental Botany. 59(8): 2133–2146.

21. Li H, Durbin R. 2009. Fast and accurate short read alignment with Burrows–Wheeler transform. Bioinformatics. 25(14): 1754–1760.

22. Li H, Handsaker B, Wysoker A, Fennell T, Ruan J, Homer N, Marth G, Abecasis G, Durbin R, 1000 Genome Project Data Processing Subgroup. 2009. The Sequence Alignment/Map format and SAMtools. Bioinformatics. 25(16): 2078–2079.

23. Li M, He P, Zhao Z, Liu J, Liu H, Ma S, Shen Y, Li B. 2023. Effect of temperature on betacyanins synthesis and the transcriptome of Suaeda salsa. Front Plant Sci. 14: 1203089.

24. Li X, Li C, Zhu J, Zhong S, Zhu H, Zhang X. 2023. Functions and mechanisms of RNA helicases in plants. Journal of Experimental Botany, 74(7): 2295–2310.

25. Liao Y, Smyth GK, Shi W. 2019. The R package Rsubread is easier, faster, cheaper and better for alignment and quantification of RNA sequencing reads, Nucleic Acids Research, 47(8): e47.

26. Lin Y-P, Wu T-H, Chan Y-K, van Zonneveld M, Schafleitner R. 2021. *De novo* SNP calling reveals the candidate genes regulating days to flowering through interspecies GWAS of Amaranthus genus. The Plant Genome. 15: e20206.

27. Liu S, Zheng X, Pan J, Peng L, Cheng C, Wang X, Zhao C, Zhang Z, Lin Y, XuHan X et al. 2019. RNA-sequencing analysis reveals betalains metabolism in the leaf of *Amaranthus tricolor* L. PLOS ONE. 14(4): e0216001.

28. Love MI, Huber W, Anders S. 2014. Moderated estimation of fold change and dispersion for RNA-seq data with DESeq2. Genome Biology. 15(12): 550.

29. Marçais G, Kingsford C. 2011. A fast, lock-free approach for efficient parallel counting of occurrences of *k*-mers. Bioinformatics. 27: 764–770.

30. McKenna A, Hanna M, Banks E, Sivachenko A, Cibulskis K, Kernytsky A, Garimella K, Altshuler D, Gabriel S, Daly M et al. 2010. The Genome Analysis Toolkit: a MapReduce framework for analyzing next-generation DNA sequencing data. Genome Research. 20(9): 1297–1303.

31. Singh N, Samarth RM, Vashishth A, Pareek A. 2024. Amaranthus as a potential dietary supplement in sports nutrition. CyTA - Journal of Food 22:1.

32. Nguyen DC, Tran DS, Tran TTH, Ohsawa R, Yoshioka Y. 2019. Genetic diversity of leafy amaranth (*Amaranthus tricolor* L.) resources in Vietnam. Breeding Science. 69(4): 640–650.

33. Pertea M, Pertea GM, Antonescu CM, Chang TC, Mendell JT, Salzberg SL. 2015. StringTie enables improved reconstruction of a transcriptome from RNA-seq reads. Nature Biotechnology. 33(3): 290–295.

34. Petroni K, Tonelli C. 2011. Recent advances on the regulation of anthocyanin synthesis in reproductive organs. Plant Science, 181(3): 219–229.

35. Rastas P. 2017. Lep-MAP3: robust linkage mapping even for low-coverage whole genome sequencing data. Bioinformatics. 33(23): 3726–3732.

36. Ravichandran K, Saw NMMT, Mohdaly AAA, Gabr AMM, Kastell A, Riedel H, Cai Z, Knorr D, Smetanska I. 2013. Impact of processing of red beet on betalain content and antioxidant activity. Food Research International. 50(2): 670–675.

37. Reyes-Rosales A, Cabrales-Orona G, Martínez-Gallardo NA, Sánchez-Segura L, Padilla-Escamilla JP, Palmeros-Suárez PA, Délano-Frier JP. 2023. Identification of genetic and biochemical mechanisms associated with heat shock and heat stress adaptation in grain amaranths. Frontiers in Plant Science. 14: 1101375.

38. Sarker U, Lin YP, Oba S, Yoshioka Y, Hoshikawa K. 2022. Prospects and potentials of underutilized leafy Amaranths as vegetable use for health-promotion. Plant Physiology and Biochemistry: PPB. 182: 104–123.

39. Shirasawa K, Hirakawa H, Isobe S. 2016. Analytical workflow of double-digest restriction site-associated DNA sequencing based on empirical and in silico optimization in tomato. DNA Research: an International Journal for Rapid Publication of Reports on Genes and Genomes. 23(2): 145–153.

40. Smit A, Hubley R, Green P. 2015. RepeatMasker Open-4.0. 2013–2015. https://github.com/rmhubley/RepeatMasker

41. Stanke M, Diekhans M, Baertsch R, Haussler D. 2008. Using native and syntenically mapped cDNA alignments to improve de novo gene finding. Bioinformatics. 24(5): 637–644.

42. Tang H, Zhang X, Miao C, Zhang J, Ming R, Schnable JC, Schnable PS, Lyons E, Lu J. 2015. ALLMAPS: robust scaffold ordering based on multiple maps. Genome Biology. 16(1): 3.

43. Wang J, Zhang Z. 2021. GAPIT version 3: boosting power and accuracy for genomic association and prediction. Genomics, Proteomics & Bioinformatics. 19(4): 629–640.

44. Wickham H. 2016. ggplot2: Elegant Graphics for Data Analysis. Springer-Verlag New York. ISBN 978-3-319-24277-4.

45. Yang R, Huang T, Song W, An Z, Lai Z, Liu S. 2024. Identification of WRKY gene family members in amaranth based on a transcriptome database and functional analysis of *AtrWRKY42-2* in betalain metabolism. Frontiers in Plant Science, 14, 1300522.

46. Xiao F, Zheng Y, Chen J, Zhao C, Chen H, Wang L, Liu S. 2021. Selection and validation of reference genes in all-red Amaranth (*Amaranthus tricolor* L.) seedlings under different culture conditions. The Journal of Horticultural Science and Biotechnology, 96(5): 604–613.

47. Xie F, Chen C, Chen J, Chen J, Hua Q, Shah K, Zhang Z, Zhao J, Hu G, Chen J, Qin Y. 2023. Betalain biosynthesis in red pulp pitaya is regulated via HuMYB132: a R-R type MYB transcription factor. BMC Plant Biol 23, 28.

48. Xue Y, Li K, Feng W, Lai Z, Liu S. 2025. Identification of R2R3-MYB Transcription Factor Family Based on *Amaranthus tricolor* Genome and *AtrMYB72* Promoting Betalain Biosynthesis by Directly Activating *AtrCYP76AD1* Expression. Plants. 14(3):324.

49. Yarra R, Xue Y. 2020. Ectopic expression of nucleolar DEAD-Box RNA helicase *OsTOGR1* confers improved heat stress tolerance in transgenic Chinese cabbage. Plant Cell Reports. 39(12): 1803–1814.

50. Yin L, Zhang H, Tang Z, Xu J, Yin D, Zhang Z, Yuan X, Zhu M, Zhao S, Li X, Liu X. 2021. rMVP: A Memory-efficient, Visualization-enhanced, and Parallel-accelerated tool for Genome-Wide Association Study, Genomics, Proteomics & Bioinformatics. 19(4), 619–628

51. Zhang L, Chen C, Xie F, Hua Q, Zhang Z, Zhang R, Chen J, Zhao J, Hu G, Qin Y. 2020. A Novel WRKY Transcription Factor HmoWRKY40 Associated with Betalain Biosynthesis in Pitaya (*Hylocereus monacanthus*) through Regulating HmoCYP76AD1. International Journal of Molecular Sciences. 22(4): 2171.

