## Supplemental Figures for "Integrated GWAS and transcriptome analyses reveal candidate genes associated with betalain-related pigmentation under heat stress in *Amaranthus tricolor*"

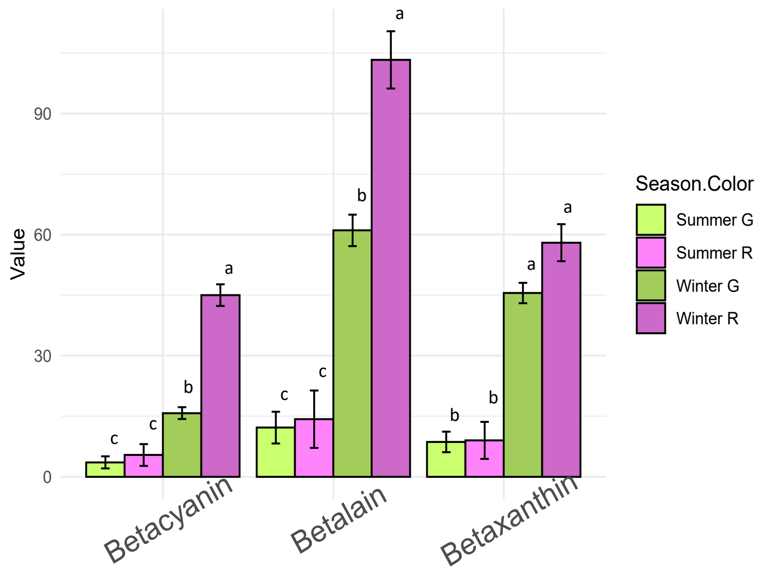


**Supplementary Figure 1 Comparisons of betalain between seasons and leaf colors.** The error bars indicate the standard errors. The letters upon the bars refer to the statistical groups resulting from the Tukey's HSD test.

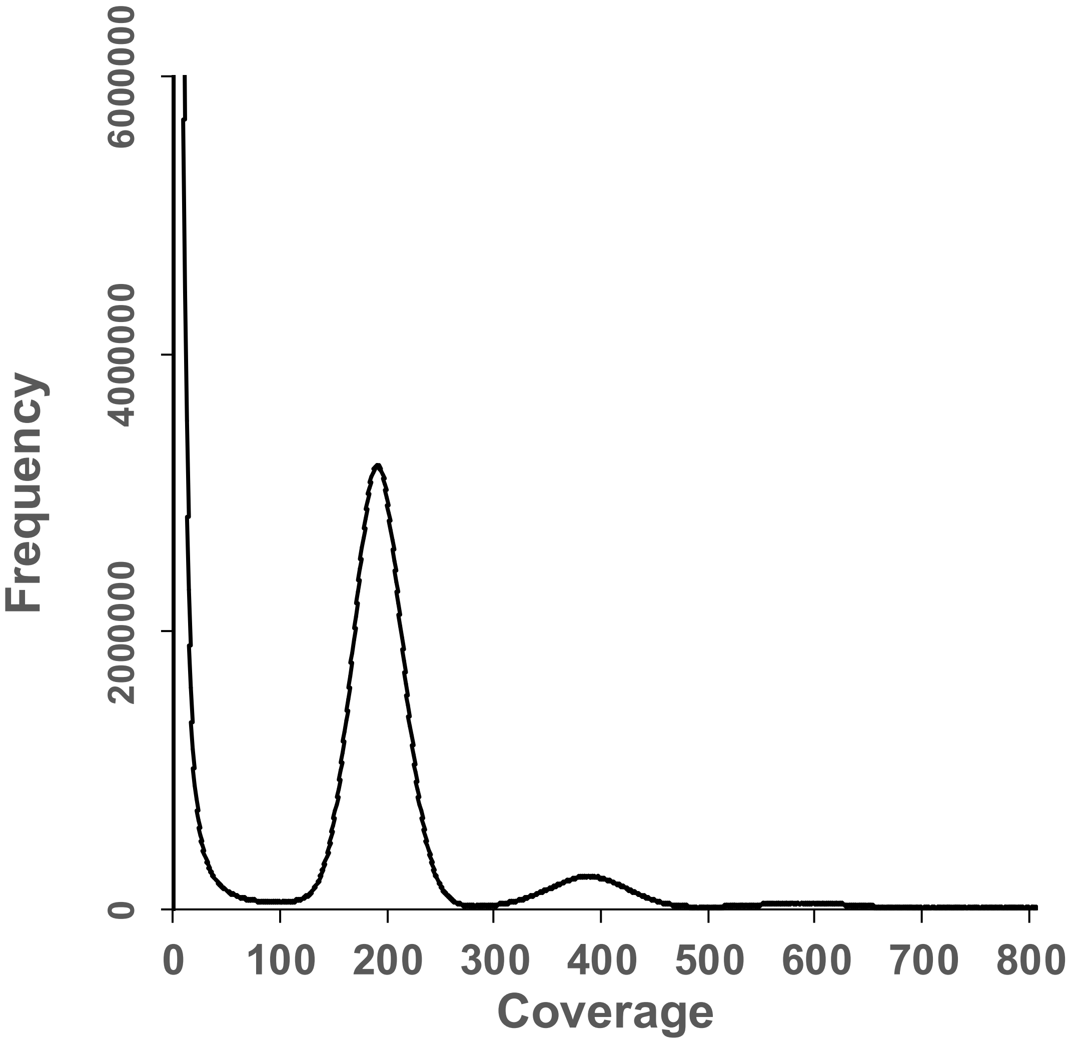


**Supplementary Figure 2 The genome size of Bayam.** *K*-mer analysis indicates the genome size of Bayam is about 596.4 Mb

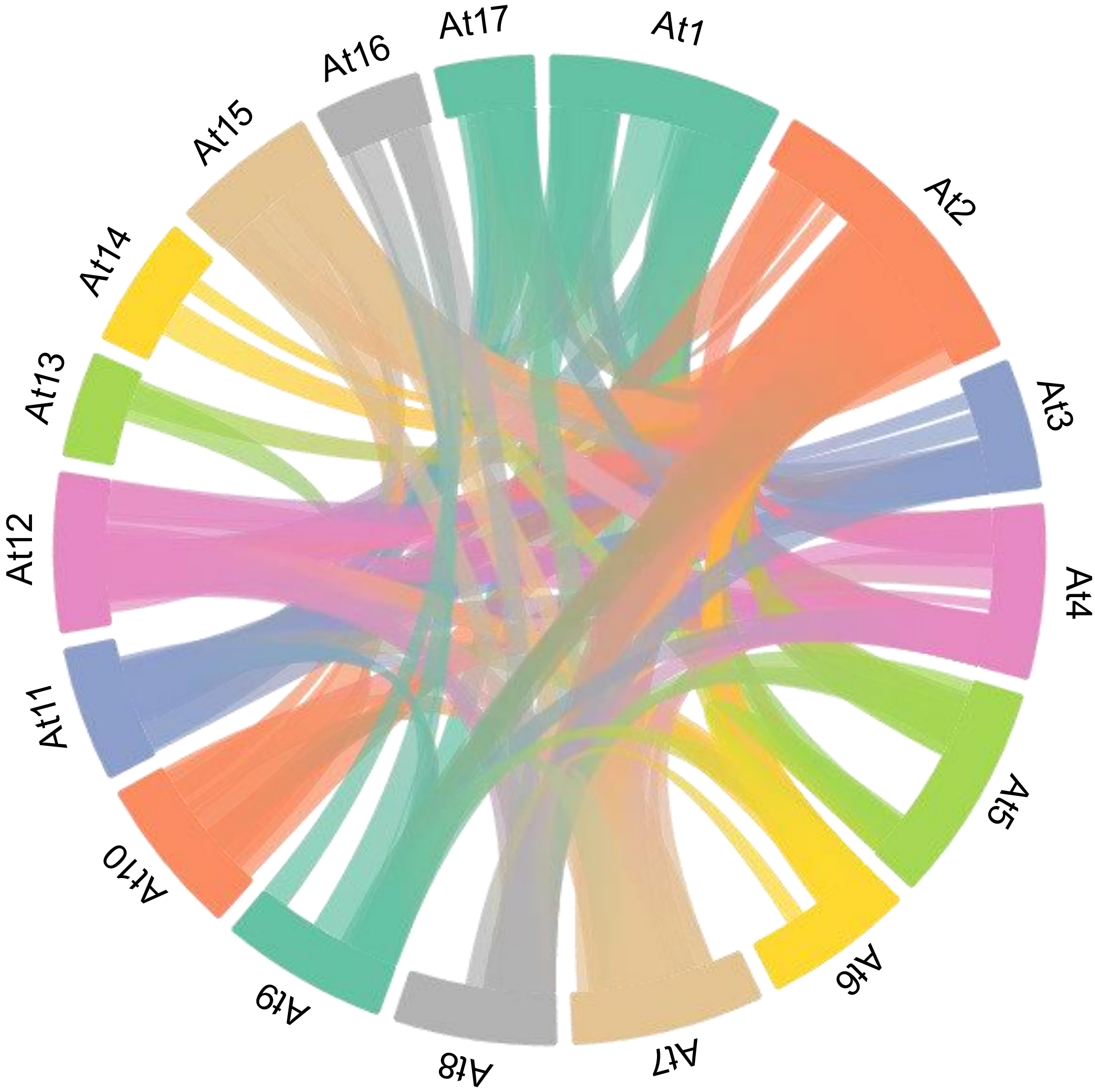


**Supplementary Figure 3 Genome duplication in the *A. tricolor* cv. Bayam.** Similar genome segments are connected with lines.

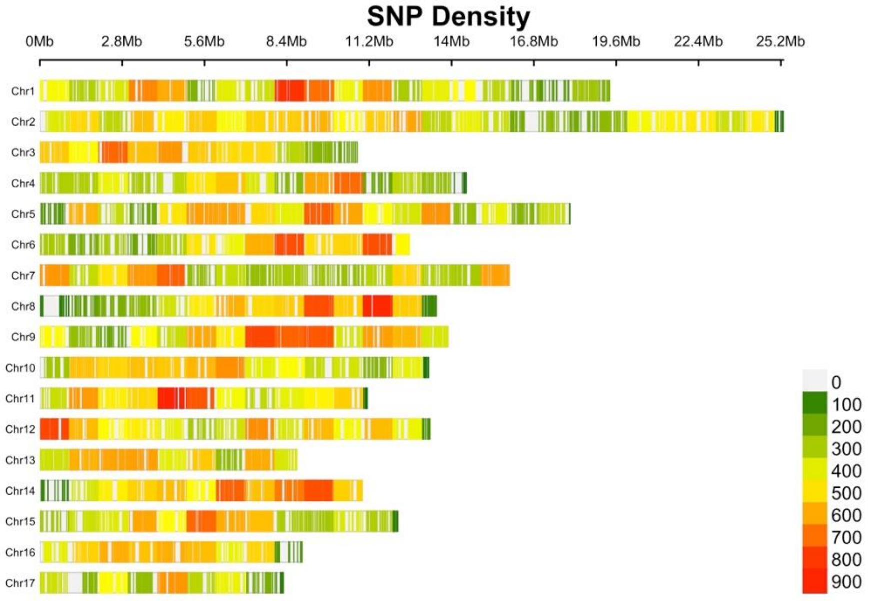


**Supplementary Figure 4 Visualization of SNP Distribution.** This figure illustrates the distribution of single nucleotide polymorphisms (SNPs) across 17 chromosomes. The horizontal axis represents chromosomal positions in megabases (Mb), while the vertical axis lists chromosome numbers. Color intensity indicates SNP density, ranging from green (low density) to red (high density), as shown in the legend (0–900). Regions in red correspond to high SNP concentration, whereas green and yellow areas indicate lower density. This distribution pattern provides valuable insights for genetic diversity analysis, quantitative trait locus (QTL) mapping, and genomic selection.

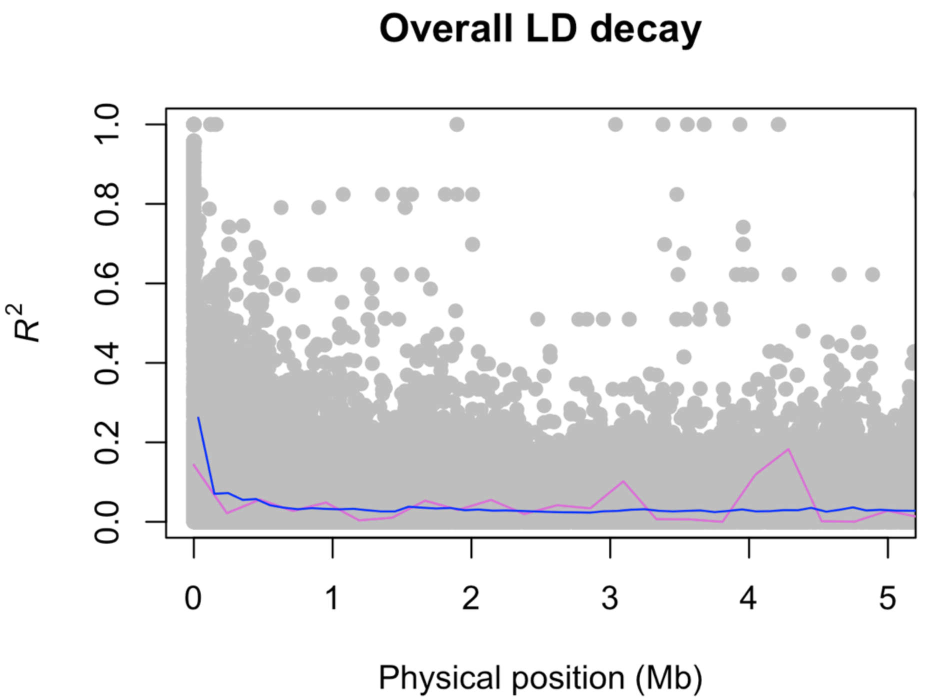


**Supplementary Figure 5 Genome-wide linkage disequilibrium (LD) decay pattern.** The plot illustrates the decline of LD with increasing physical distance across the genome. The x-axis represents physical distance in megabases (Mb), and the y-axis shows the pairwise LD measure (*R*^2) between SNPs. Gray dots indicate individual SNP pairs, while the purple line represents the sliding window average and the blue line shows the LOESS-smoothed LD decay curve. LD decreases rapidly within short distances and stabilizes beyond several hundred kilobases, reflecting the extent of recombination and genetic diversity in the population.

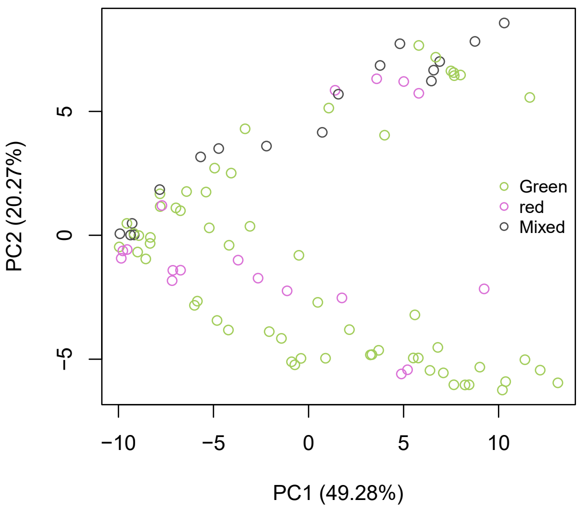


**Supplementary Figure 6 Principal Coordinate Analysis of the core collection.** Colors indicate the leaf color of each accession.

(A)

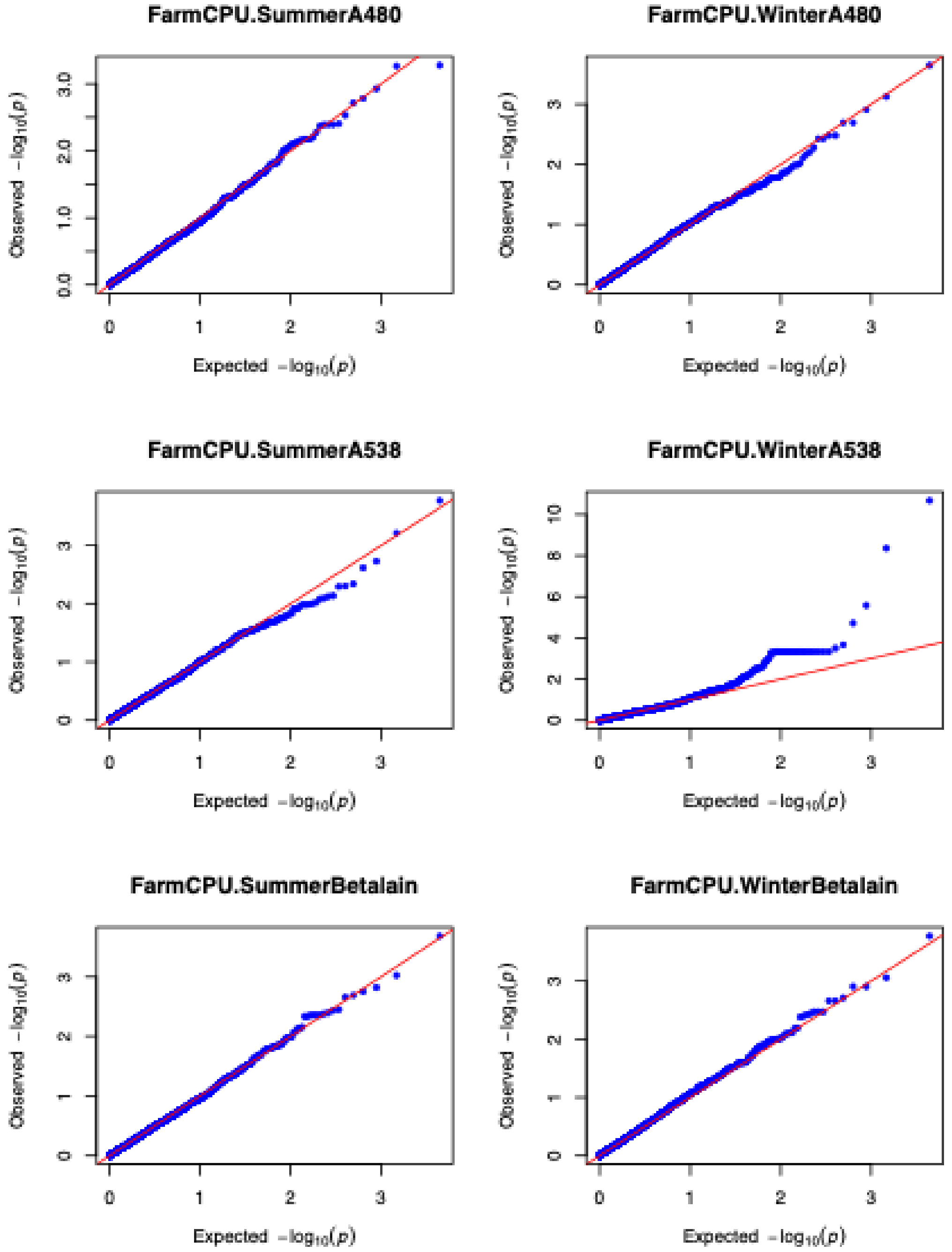


**Supplementary Figure 7 Q-Q plots of betaxanthin-dominated (A480), betacyanin-dominant (A538), and betalain absorbance index (A480 + A538).** A) and B) refer to the Q-Q plots of the FarmCPU and the Blink models, respectively.

(B)

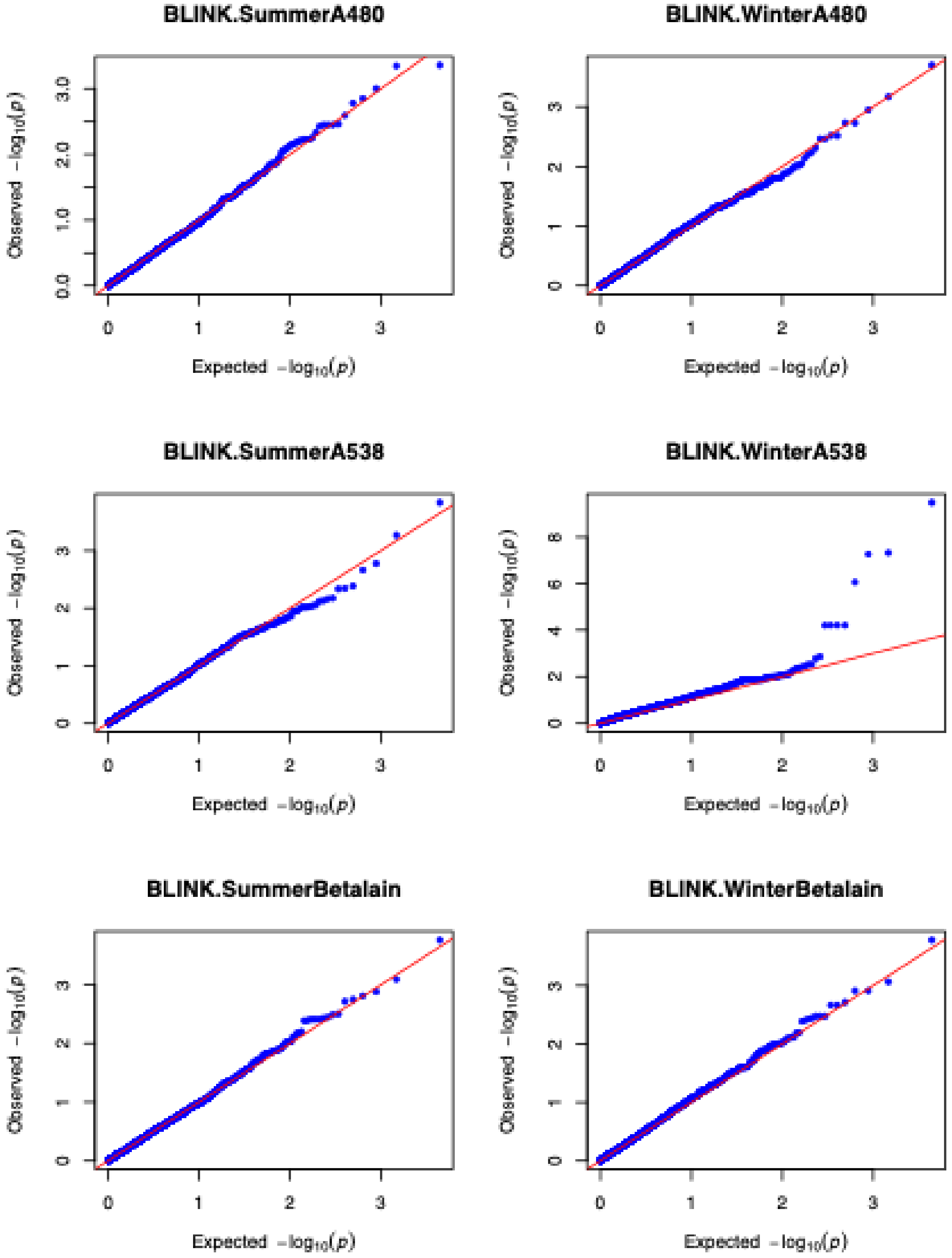


**Supplementary Figure 7 (continued) Q-Q plots of betaxanthin-dominated (A480), betacyanin-dominant (A538), and betalain absorbance index (A480 + A538).** A) and B) refer to the Q-Q plots of the FarmCPU and the Blink models, respectively.

(A)

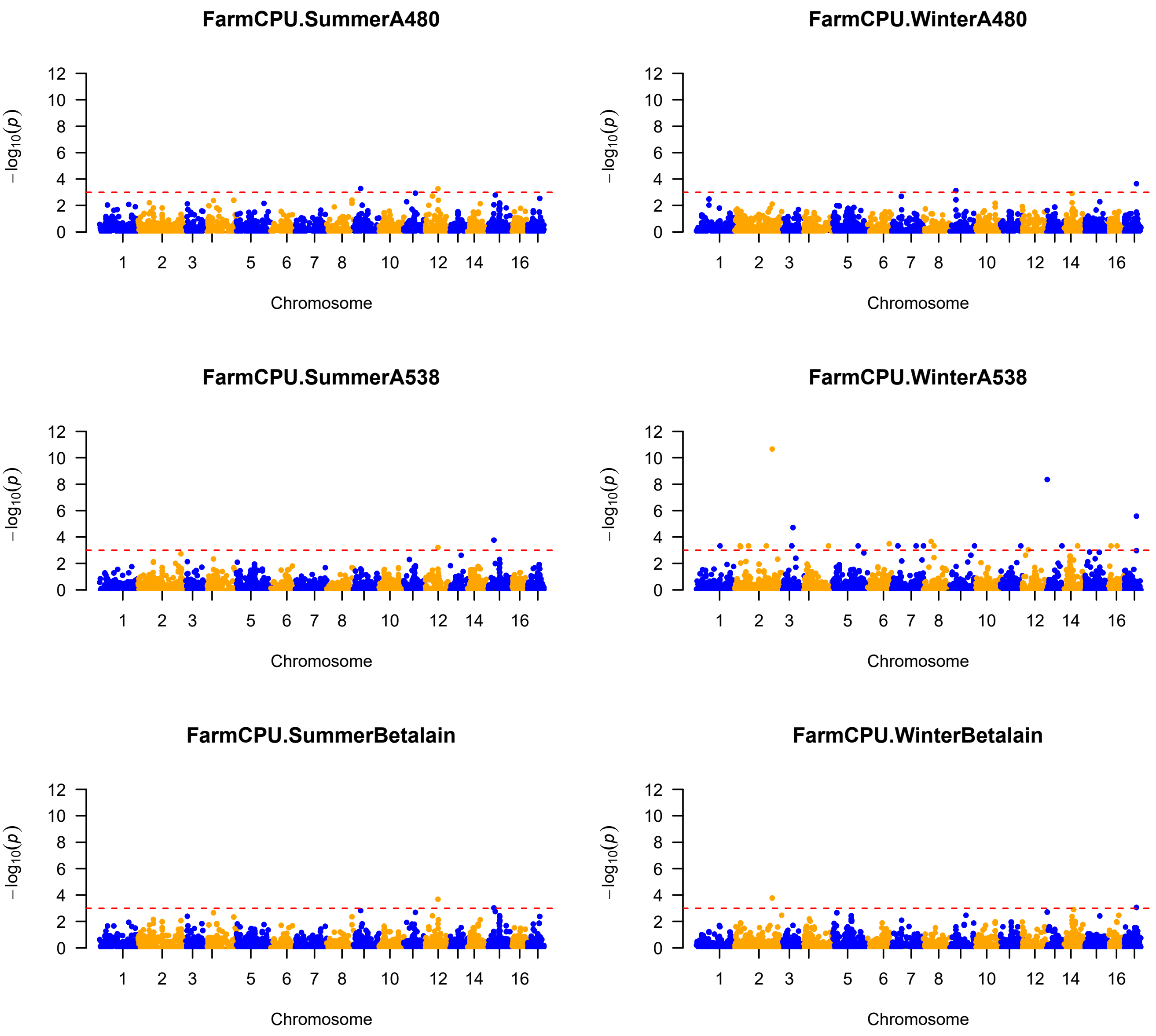


**Supplementary Figure 8 Manhattan plots of betaxanthin-dominated (A480), betacyanin-dominant (A538), and betalain absorbance index (A480 + A538).** A) The Manhattan plots of the FarmCPU model. B) The Manhattan plots of the Blink model. The horizontal red dashed line indicates the significance cutoff (*p value* < 0.001).

(B)

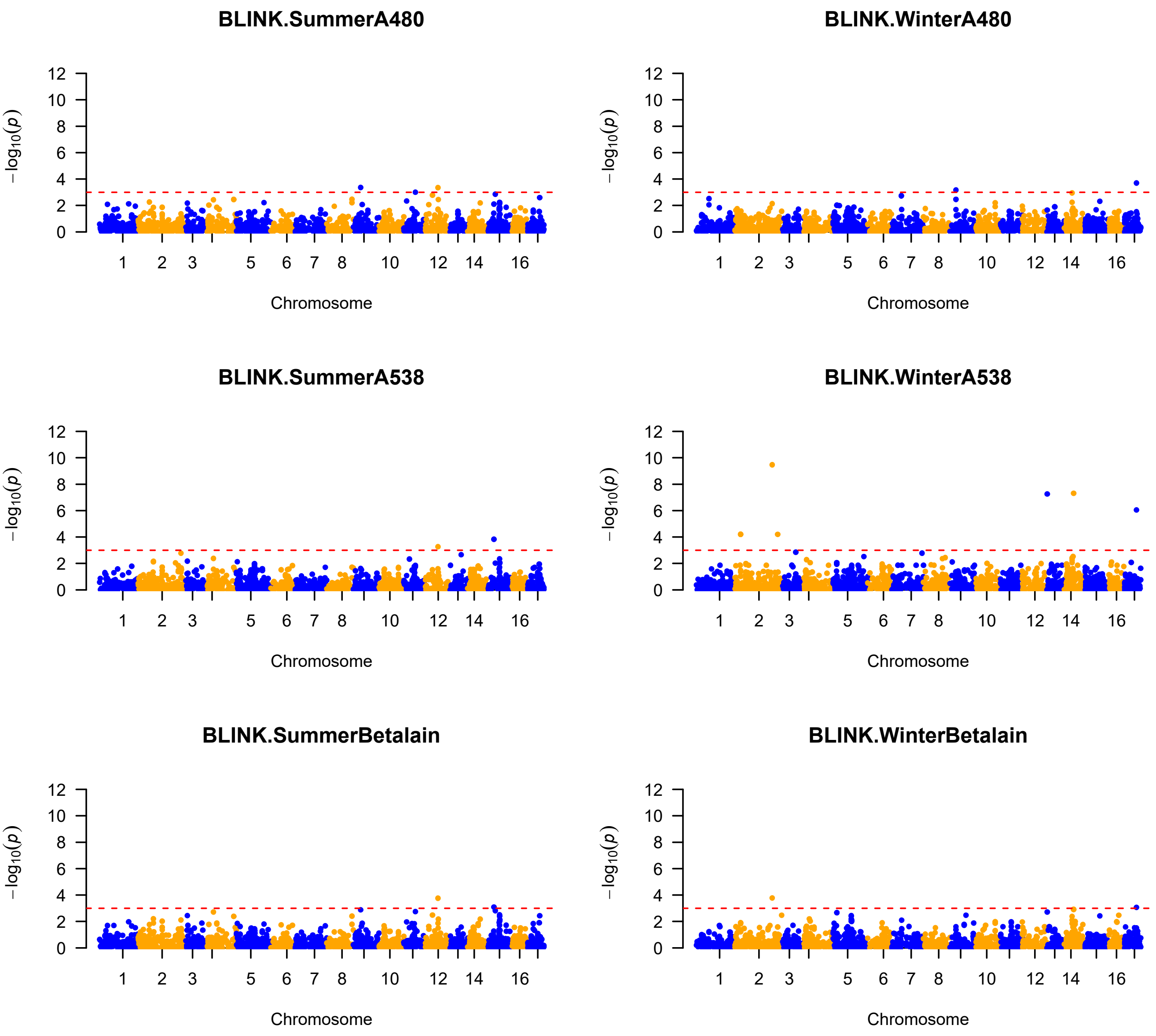


**Supplementary Figure 8 (continued) Manhattan plots of betaxanthin-dominated (A480), betacyanin-dominant (A538), and betalain absorbance index (A480 + A538).** A) The Manhattan plots of the FarmCPU model. B) The Manhattan plots of the Blink model. The horizontal red dashed line indicates the significance cutoff (*p value < 0.001*).

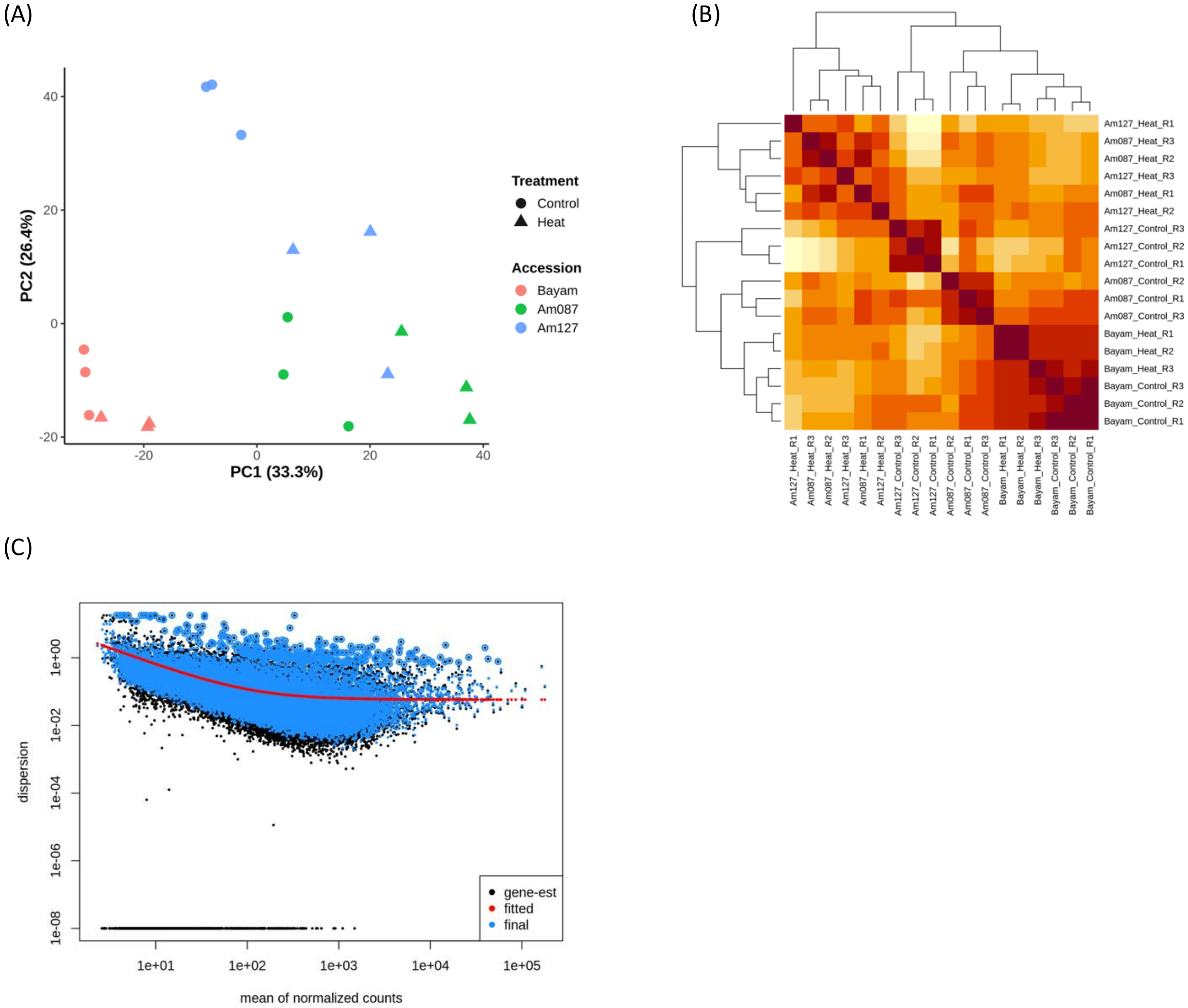


**Supplementary Figure 9 RNA-seq sample-level and DESeq2 model diagnostics.** (A) Principal component analysis of the 18 RNA-seq libraries based on variance-stabilized expression values of the 500 most variable genes. Colors indicate accession and symbols indicate treatment. PC1 and PC2 explain 33.3% and 26.4% of the total variance, respectively. (B) Sample-to-sample Pearson correlation heatmap calculated from variance-stabilized expression values across the 18 libraries. (C) DESeq2 dispersion diagnostic showing gene-wise dispersion estimates and the fitted mean–dispersion relationship.

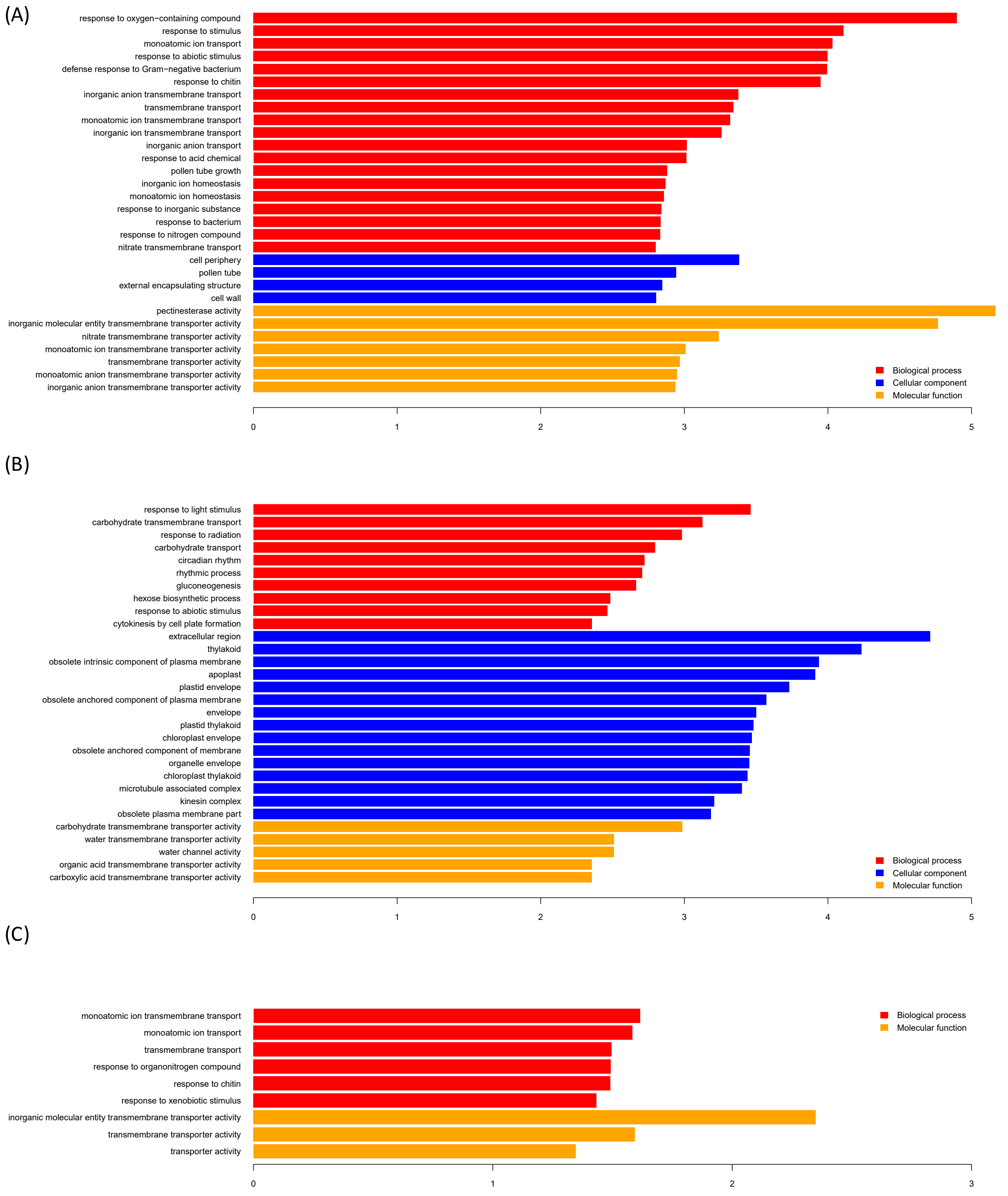


**Supplementary Figure 10 GO enrichment analyses.** A) and B) are the top 30 enriched GO in Am087 and Am127, respectively. C) is the enriched GO shared by Am087 and Am127.

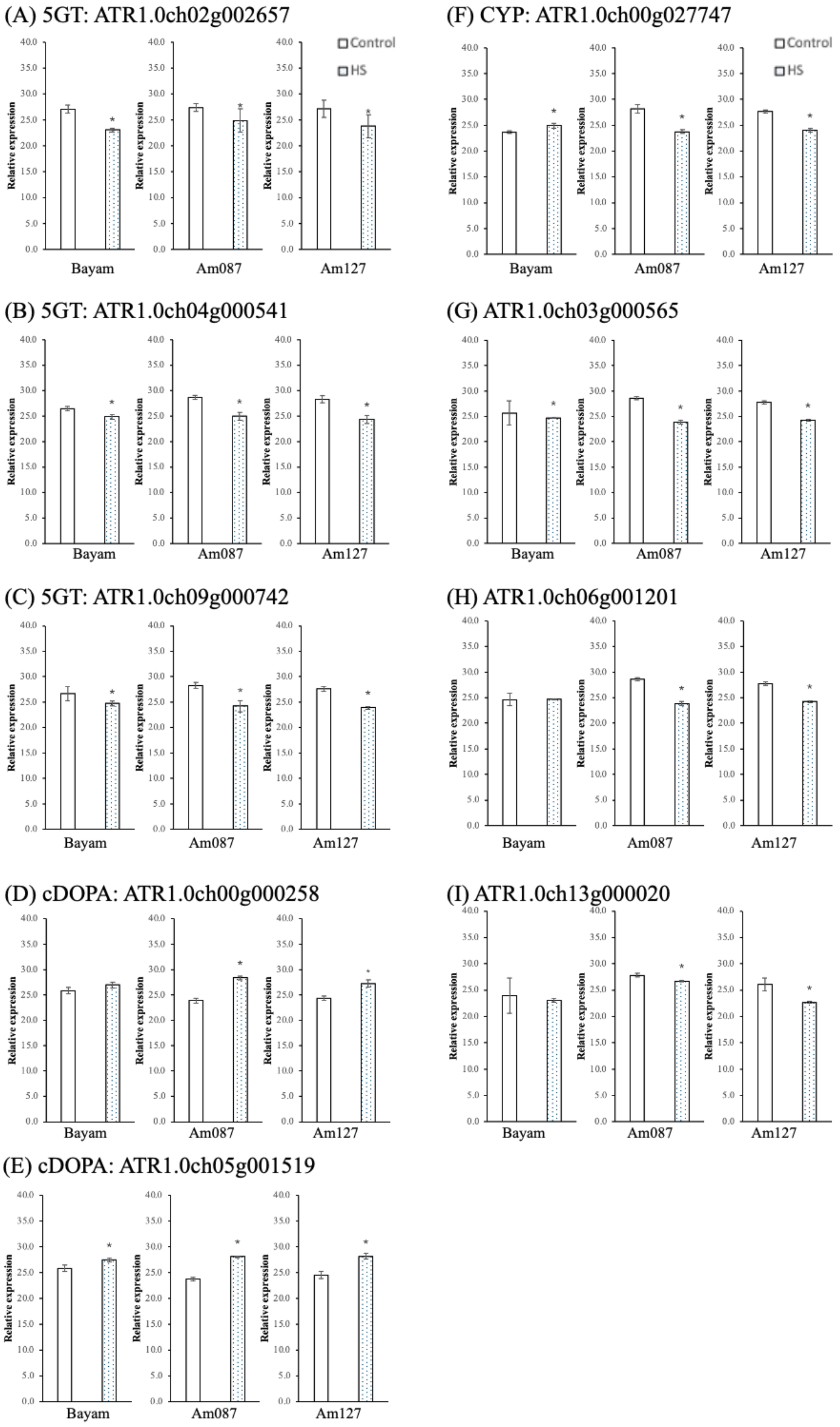


**Supplementary Figure 11 Relative expression patterns of candidate genes in different *Amaranthus* accessions under control and heat stress conditions.** (A) 5GT: *ATR1.0ch02g002657*, (B) 5GT: *ATR1.0ch04g000541*, (C) 5GT: *ATR1.0ch09g000742*, (D) cDOPA: *ATR1.0ch00g000258*, (E) cDOPA: *ATR1.0ch05g001519*, (F) CYP: *ATR1.0ch00g027747*, (G) *ATR1.0ch03g000565*, (H) *ATR1.0ch06g001201*, (I) *ATR1.0ch13g000020*. Relative expression levels were measured by qRT-PCR in leaves of *Amaranthus* accessions: Bayam, Am087, and Am127. The heat stress condition was as described in Materials and Methods. Gene expression was compared across three *Amaranthus* accessions: Bayam, Am087, and Am127. The bars represent the mean values (n = 3) and intervals over the bars indicate the standard deviation of the means. * Significant change (*p value* $< 0.05$) relative to the control.
